# Transmembrane signalling by a bionic receptor: biological input and output, chemical mechanism of signal transduction

**DOI:** 10.1101/2021.07.25.453684

**Authors:** Kaja B. Løvschall, Pere Monge, Line F. Nielsen, Sandra Stevanovic, Raoul Walther, Alexander N. Zelikin

## Abstract

Signal transduction through sealed biological membranes is among the most important evolutionary achievements. Herein, we focus on the development of artificial signal transduction mechanisms and engineer a bionic receptor with capacity of transduction of biological signals across biological membranes using tools of chemistry. The bionic receptor described in this work exhibits similarity with the natural counterpart in the most essential characteristics: in having an exofacial ligand for signal capture, in being membrane anchored, and in featuring a releasable secondary messenger molecule, which performs enzyme activation in the endo volume. The main difference with the natural receptors is that signal transduction across the lipid bilayer was performed using the tools of organic chemistry, namely a self-immolative linker. The highest novelty of our work is that the artificial signalling cascade designed herein achieved transmembrane activation of enzymatic activity, as is the hallmark of activity by natural signalling receptors.

---

Mechanisms of transfer of information across the cell membrane lipid bilayer are among the most important evolutionary adaptations which allow cells to sense external environment and to communicate within multicellular ensembles. From the simplest to the most advanced cells, these mechanisms rely on transmembrane proteins, termed signalling receptors, which perceive information at the cell surface and communicate it to the cell interior. ^1–3^ With the exception of ion channels, transfer of chemical information by receptors is performed through sealed membranes, without compromising the integrity of the lipid bilayer. Performance of the transmembrane receptor proteins is robust and elegant at the same time. Modern science can successfully reproduce this activity using chimeric receptors, ^4,5 6^ whereas the design of bionic receptors remains largely unfulfilled. As such, receptor mimicry and transfer of information through sealed biomolecular membranes using non-natural tools is a grand fundamental challenge with only few successes.

One class of bionic signalling receptors, initially designed by Hunter and Williams et al ^7 8^ and later adopted by Schrader et al ^9^, relies on the toolbox of chemically induced dimerization and uses membrane-spanning cholesterol dimers. The dimerization event at the exo-surface (due to e.g. oxidation of thiols into a disulfide, ^7^ Cu^2+^ bridging of two monovalent ligands ^8^ etc.) evoked proximity of the two transmembrane molecules and ensued dimerization of endofacial termini of these molecules, without compromising integrity of the lipid bilayer. Typical results in these studies included a release of a UV-active molecule^7^ or an energy transfer event between the dimerizing “sub-units” ^9^. Another class of bionic receptors was designed in recent years by Hunter et al ^10–12^ using molecules that exhibit controlled “bobber”-like translocation across the lipid bilayer. In these cases, receptor activation (by a change in solution pH ^10^, the presence of copper ions,^11^ or a competitive ligand displacement event ^12^) ensued a cross-membrane movement of the receptor molecule, which resulted in an exposure of a Zn-coordinating ligand as a metalloenzyme mimic. Finally, Clayden et al ^13^ developed peptide foldamers that exhibit in-membrane conformational change, mimicking performance of natural receptors, in response to an enkephalin agonist. Each of these results is highly important, but the field is still at its infancy and existing examples of bionic signalling receptors have limited biological relevance in both receptor activation mechanism and the receptor output.

In this work, we take a major step forward and engineer a bionic receptor which exhibits similarity to the natural counterpart in the most essential characteristics, namely in having an exofacial ligand to capture a biological input, in being membrane anchored, and in featuring a releasable, biological secondary messenger molecule, which performs downstream enzyme activation (Figure 1A). The main difference is that signal transduction across the lipid bilayer is performed herein using tools of organic chemistry rather than protein molecules.

**Figure 1.**
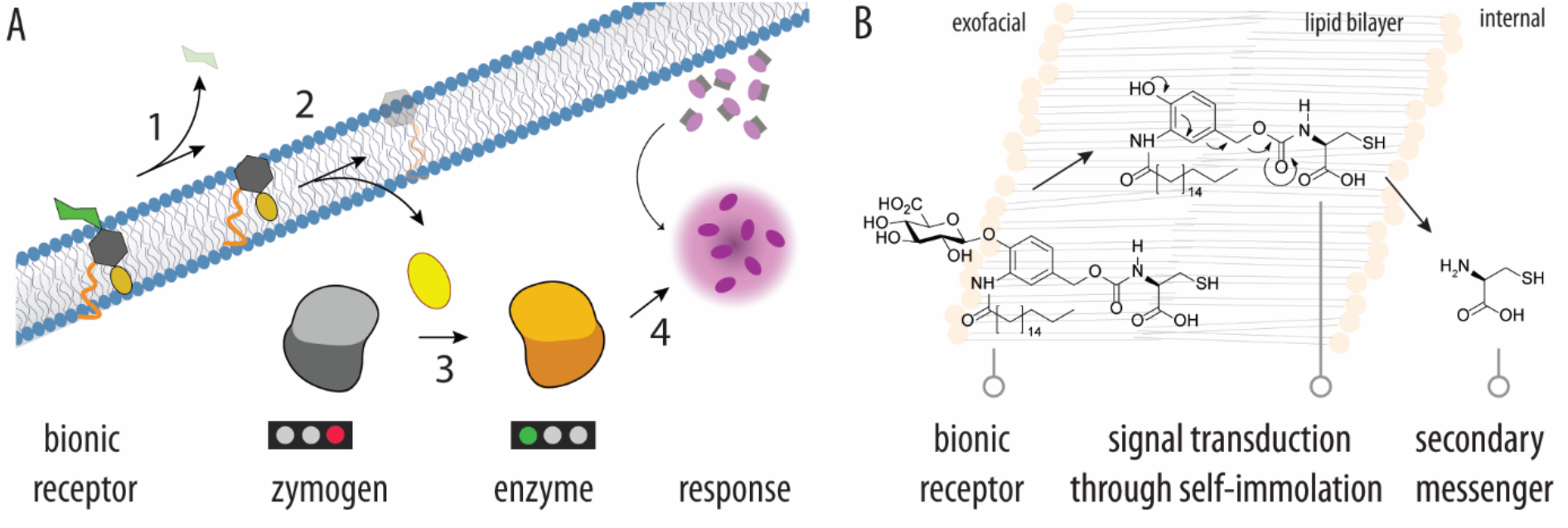
**(A) Schematic illustration of the proposed concept of transmembrane signaling using a lipid-bilayer anchored signal transduction molecule:** 1) receptor activation at the exofacial side of the membrane, 2) self-immolation and ensuing generation of the secondary messenger which enters the inner compartment; 3) activation of the zymogen by the secondary messenger; 4) secondary signaling events or signal amplification via enzymatic catalysis. **(B) Specific chemistry realized in this work as transmembrane signaling**: artificial receptor features an exofacial glucuronic acid moiety for receptor activation using the corresponding enzyme (GUS), lipid-modified *p*-hydroxybenzyl alcohol is a self-immolative linker which acts as the chemical mechanism of signal transduction, natural amino acid L-cysteine is released and acts a secondary messenger.

The bionic receptor designed in this work features a self-immolative molecule as the mechanism of signal transduction. Self-immolative linkers (SIL) such as *p*-hydroxybenzyl alcohol are highly successful in the design of prodrugs, both in academia and in industry.^14–19^ These cunning tools were initially developed to decrease steric hindrance around the scissile bond and thus enhance its accessibility to an enzyme-activator. The latter acts to remove the prodrug masking group, and this initiates an electronic cascade which ultimately leads to the traceless release of the pristine drug. In our recent work, we demonstrated that SIL performance and drug release were efficient even when prodrugs were anchored at the mammalian cell surface, in which case the prodrugs comprised artificial receptors for triggered deactivation of the engineered cells.^20^

These results led us to recognize that SIL can serve as a bionic mechanism for signal transduction across sealed biological membranes, thus addressing the highest challenge in receptor mimicry. Below, we realize this opportunity synthetically. The highest novelty of our work is that artificial signalling cascade designed in this work using SIL achieved transmembrane activation of enzymatic catalysis, as is the hallmark of activity by natural signalling receptors.

The receptor molecule proposed herein is an amphiphile, engineered on the *p*-hydroxybenzyl alcohol scaffold. It contains a highly polar glucuronic acid as an exofacial receptor masking group, which engages with an incoming trigger to activate signal transduction (Figure 1B). Glucuronides are substrates to a natural enzyme, β-glucuronidase (GUS), which was chosen to serve as the receptor’s cognate binding partner and activating biomolecule. Transmembrane and/or downstream signalling events in nature most commonly rely on enzyme (de)activation. ^2,3^ To our knowledge, this opportunity has never been realized using bionic receptors, and it was chosen in this work as the prime design challenge. To this end, we focused on a cysteinyl protease, papain. Activity of this enzyme is blocked by chemical modifications of the thiol group in the active site, for example by conversion into a disulfide functionality.^21^ Papain catalytic activity is restored via thiol-disulfide exchange with another thiol-containing molecule – thus presenting an opportunity to achieve chemically triggered activation of enzymatic catalysis. This notion suggested the final composition of the artificial receptor molecule, specifically that the secondary messenger released by the receptor upon exofacial activation must be a thiol-containing solute. To complete biological relevance of the bionic receptor designed herein, we chose to use L-cysteine as a natural thiol-containing molecule (Figure 1B).

The synthetic path to the enzyme-activating artificial receptor (EAR) molecule consisted of an 11-step synthesis, starting with methyl 1,2,3,4-tetra-O-acetyl-β-D-glucuronate(Figure 2A). 4-hydroxy-3-nitro-benzaldehyde formed the template of the SIL scaffold, chemical *O*-glycosylation delivered the required beta-diastereomer for enzyme recognition, and reduction of the nitro- and aldehyde moieties delivered the chemical handles for the lipid bilayer anchoring ligand and the secondary messenger, respectively. For bilayer anchoring, we used stearic acid, a natural fatty acid. The acyl chloride reacted smoothly with the amine in the presence of triethylamine to deliver the desired amide **7**. *S*-Trityl-L-cysteine was introduced via carbonate exchange reaction with mixed carbonate **9** to afford **10** in 80% yield. Finally, deprotection of glucuronic acid afforded the *S*-trityl protected EAR molecule (**11**). The latter was used in the HPLC characterization of receptor activation and release of the secondary messenger, so as to take advantage of ease of detection of the trityl- group. Compound **11** was stable in solution and revealed no signs of spontaneous decomposition. Upon addition of GUS enzyme, **11** quantitatively released *S*-trityl-L-cysteine, illustrating enzyme activated release of the (*S*-protected) secondary messenger molecule, in solution. Lastly, removal of the trityl-group using triflouroacetic acid and triisopropylsilane afforded the desired receptor molecule EAR (**12**).

**Figure 2.**
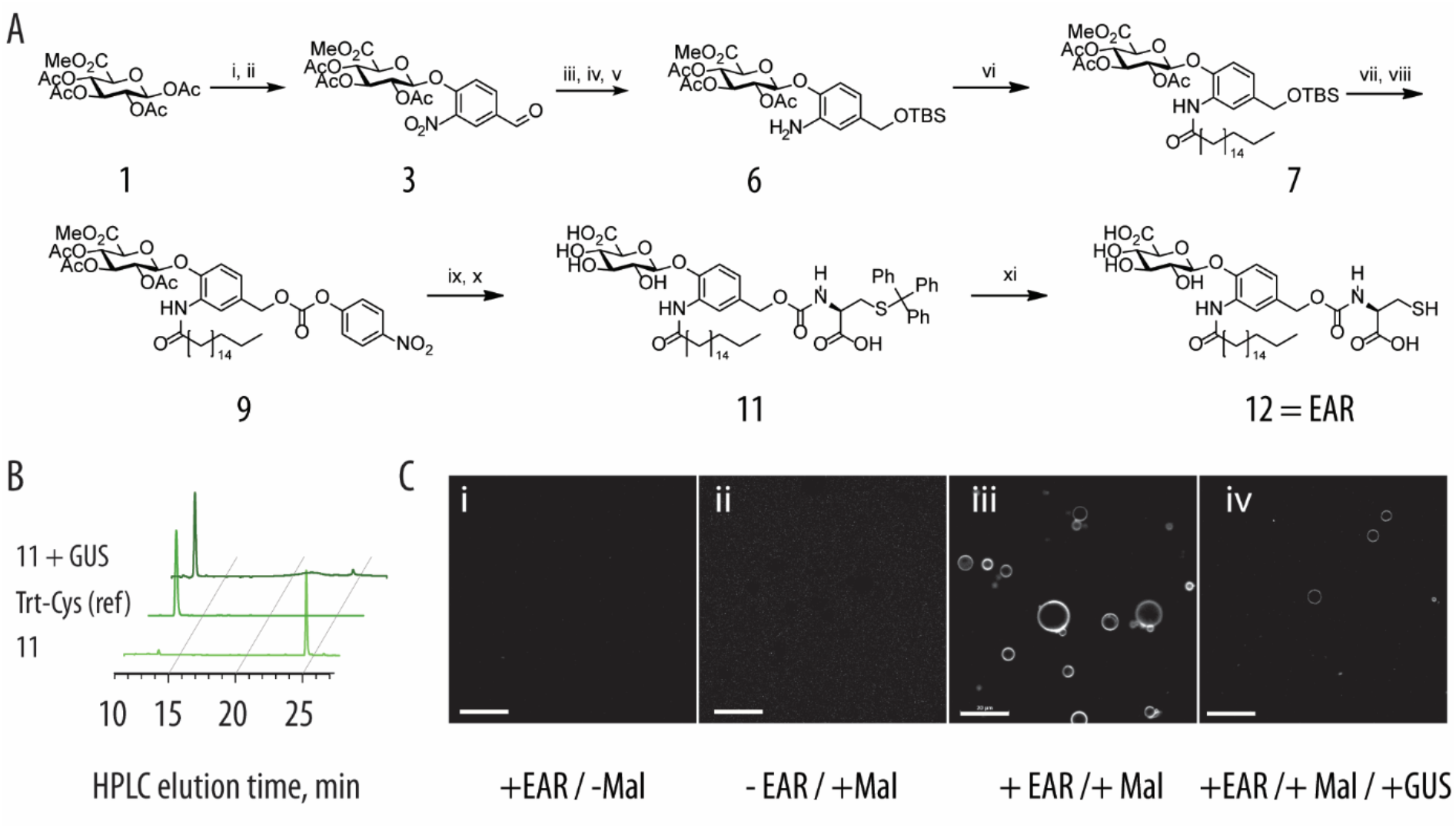
(A) Schematic illustration of the 11-step synthesis implemented to obtain the artificial receptor molecule. Conditions and reagents; i) HBr/AcOH, CH_2_Cl_2_, 0°C to r.t., 4 h, 94%, ii) 4-hydroxy-3-nitrobenzaldehyde, powdered 3Å molecular sieves, Ag_2_O, CH_3_CN, r.t., 22 h, 68%, iii) NaBH_4_, silica gel, isopropanol/CHCl_3_, 0°C, 1.5 h, 77%, iv) TBSCl, imidazole, DMAP, DMF, r.t, 41.5 h, 82%, v) Pd/C, ammonium formate, abs. EtOH, r.t., 2.5 h, 84%, vi) stearoyl chloride, TEA, CH_2_Cl_2_, 0°C to r.t., 4 .5h, 96%, vii) TEA·3HF, THF, 0°C to r.t., 42 h, 59%, viii) 4-nitrophenyl chloroformate, TEA, CH_2_Cl_2_, 0°C to r.t., 20 h, 75%, ix) *S*-Trityl-L-cysteine, TEA, CH_2_Cl_2_, 0°C to r.t., 3 h, 80%, x) NaOMe, NaOH, MeOH, 0°C to r.t., 34%, xi) TFA, (i-Pr)_3_SiH, CH_2_Cl_2_, 0°C to r.t., 2 h, 16%.; for full experimental details and compound characterization, see Supporting Information; (B) HPLC characterization of the triggered release of S-Trityl-L-cysteine upon incubation with GUS; analysis is performed on compound **11** for ease of UV-detection of the trityl-group; (C) confocal laser scanning microscopy images demonstrating lipid bilayer anchoring of EAR (compound **12**). For visualization, giant unilamelar vesicles were equipped with EAR and exposed to fluorescein maleimide (Mal). Scale bars: 20 μm.

The receptor placement and anchoring in the lipid bilayer was visualized in giant unilamelar vesicles (GUVs). GUVs were exposed to solutions of EAR (**12**) in dimethylsulfoxide and subsequently to a solution of fluorescein maleimide. Thiol-to-maleimide coupling is poised to link the fluorophore to EAR and in doing so visualize its localization. Confocal laser scanning microscopy images reveal that the EAR molecule is non-fluorescent (Figure 2Ci), and fluorescein maleimide has no inherent affinity to the lipid bilayer (Figure 2Cii). GUVs equipped with the receptor molecule and exposed to fluorescein maleimide exhibited strong fluorescence, which was confined to the lipid bilayer (Figure 2Ciii). Addition of GUS resulted in a drastic decrease of the GUV fluorescence, illustrating that receptor triggering ensues decomposition of EAR, which results in the release of cysteine and escape of the latter from the lipid bilayer (Figure 2Civ).

Encapsulation of papain zymogen into the lipid bilayer-surrounded compartments was achieved using pre-formed liposomes, and using saponin to exert temporary bilayer permeation.^22^ Zymogen activation using EAR was quantified using a fluorogenic substrate (*N*_α_-Benzoyl-L-arginine-7-amido-4-methylcoumarin) co-encapsulated with the protein, and fluorescence spectroscopy as a read-out. The EAR molecule contains a thiol functionality and receptor dormancy is conditioned by the placement of the zymogen and EAR in two different phases, liposome lumen *vs* the lipid bilayer. This separation is very efficient and in the presence of dormant EAR, the encapsulated zymogen registered only a minor “resting” signal (Figure 3A). Addition of GUS enzyme to the liposomal preparation afforded a pronounced increase in solution fluorescence. This is indicative of GUS activity on the bilayer-anchored EAR to remove the glucuronic acid masking group, ensuing decomposition of the self-immolative linker as a signal transduction mechanism, the release of the secondary messenger molecule (L-cysteine), and finally activation of the zymogen into the catalytically active papain, which decomposes the fluorogenic substrate. To our knowledge, this is the first example of a bionic receptor that connects a biological input to a biological output using tools of chemistry for transmembrane signal transduction.

**Figure 3.**
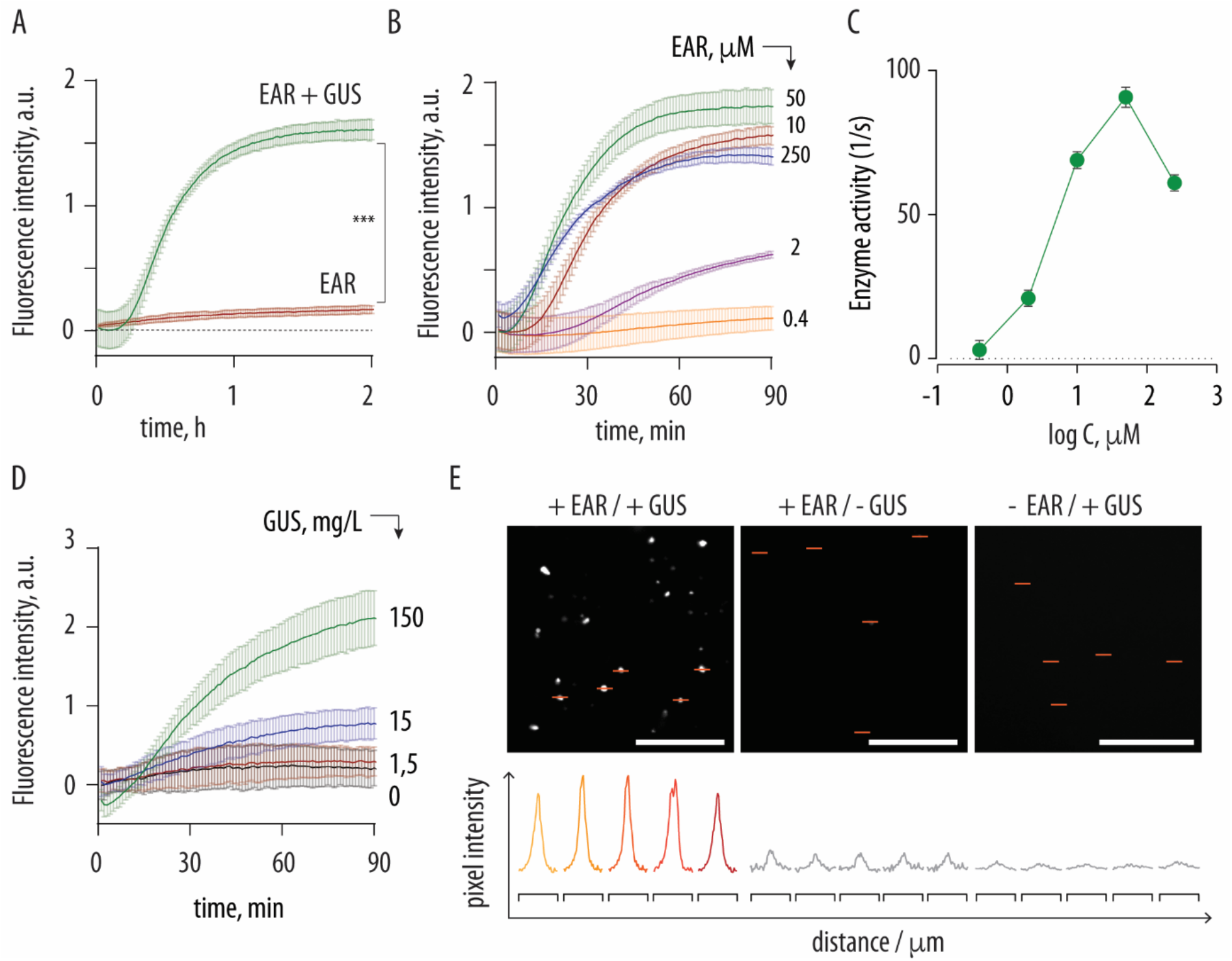
(A) proof-of-concept transmembrane activation of zymogen achieved through the performance of a bionic receptor: exofacial receptor activation by GUS initiates the signal transduction cascade which culminates in the activation of the zymogen in the interior volume, as registered by fluorescence monitoring of activity of the activated enzyme; (B) transmembrane activation of the papain zymogen in liposomes (total lipid content in solution ~2 g/L) in the presence of increasing content of EAR (from 0 to 250 μM EAR in solution); (C) rate of enzymatic reaction calculated from the data in panel B, as a function of EAR concentration; (D) transmembrane activation of the papain zymogen achieved using increasing concentration of added GUS, with activity of the activated enzyme registered on a biological substrate, FITC-casein; In panels A-D, data presented are based on at least three independent experiments and presented as mean ± SD. Statistical significance was established via an unpaired t-test *** *p* < 0.001; (E) fluorescence microscopy visualization of receptor performance in 800 nm liposomes: exofacial addition of GUS triggers signal transduction using EAR, which ensues activation of the encapsulated zymogen, as evidenced by evolution of fluorescence signal due to enzymatic decomposition of the corresponding fluorogenic substrate; zymogen activation is not registered in the absence of the enzyme and neither in the absence of receptor. Scale bars : 20 μm; intensity profile traces are 4 μm each.

With increased content of EAR per liposome, both the absolute level of attained fluorescence (Figure 3B) and the rate of enzymatic substrate conversion (Figure 3C) progressively increased and reached a maximum value at an EAR concentration of ca. 50 μM, whereas the increase to EAR content to 250 μM afforded a decrease in the rate of reaction. This observation is consistent with the value of critical micellization concentration for EAR at 90 μM, established using pyrene fluorescence.^23^ This indicates that incorporation of EAR into liposomes is favoured from the solution phase rather than from the micellar phase. At each tested concentration of EAR, the onset time for receptor performance was measured in minutes (Figure 3B), which places EAR in terms of response time between the natural G-protein coupled receptors and receptor tyrosine kinases.

To increase biological relevance of the transmembrane signal transduction cascade developed in this work, we considered that in nature, downstream signalling typically involves (de)activation of enzymes and/or protein processing. We co-encapsulated papain zymogen and its biological substrate, casein. The latter was used in a form of its derivative labelled with fluorescein to a level above self-quenching. Processing by papain should result in degradation of casein and an increase in fluorescence. Indeed, receptor activation using GUS afforded a pronounced increase in fluorescence signal, indicating successful transmembrane signalling, activation of the zymogen, and downstream signalling through the processing of a biological substrate. In these experiments, we also considered that receptor response in nature is a concentration-dependent event and occurs only if the messenger concentration is present in sufficient quantity. This was successfully realized using artificial transmembrane signalling developed herein, and casein processing was concentration dependent with regards to the amount of added GUS, Figure 3D.

Receptor performance was visualized using fluorescence microscopy and liposomes of 800 nm diameter. Zymogen-containing liposomes equipped with a dormant EAR exhibited fluorescence levels only marginally above the background level, illustrating only non-detectable activity of the receptor in its resting state (Figure 3E). Expectedly, addition of GUS to the EAR-negative liposomes afforded negligible increase in fluorescence. In stark contrast, addition of GUS enzyme to the receptor-containing liposomes led to a pronounced increase in fluorescence within liposomal compartments, viewed as sub-micrometer bright fluorescence loci, thus providing direct visualization of transmembrane signalling. Taken together, results presented in Figure 3 illustrate what is, to our knowledge, the first artificial chemical transmembrane receptor that is activated by a biological messenger, operates through generation of a biological (natural) secondary messenger molecule, and culminates in a zymogen activation event.

To validate the receptor-like mechanism of activity of EAR, we compared activation of the encapsulated zymogen by cysteine, added externally or released via the activation of EAR, at identical concentrations. In this experiment, matched efficacy of zymogen activation by the two treatments would indicate that upon receptor activation, the secondary messenger molecule is released from EAR into solution bulk, which is inconsistent with the proposed transmembrane signalling. In contrast, superior efficacy of EAR over externally administered cysteine would indicate that the secondary messenger is released from EAR directly into the host liposome. In our experiment, at L-cysteine concentration of 2 μM we observed negligible activation of the encapsulated zymogen with the soluble amino acid, Figure 4B. At the same concentration of lipids and 2 μM content of EAR, the synthesized receptor molecule produced a strong signalling event, as evidenced by pronounced evolution of solution fluorescence in response to the added GUS. This result provides the strongest evidence to postulate receptor-mediated transmembrane signalling.

**Figure 4.**
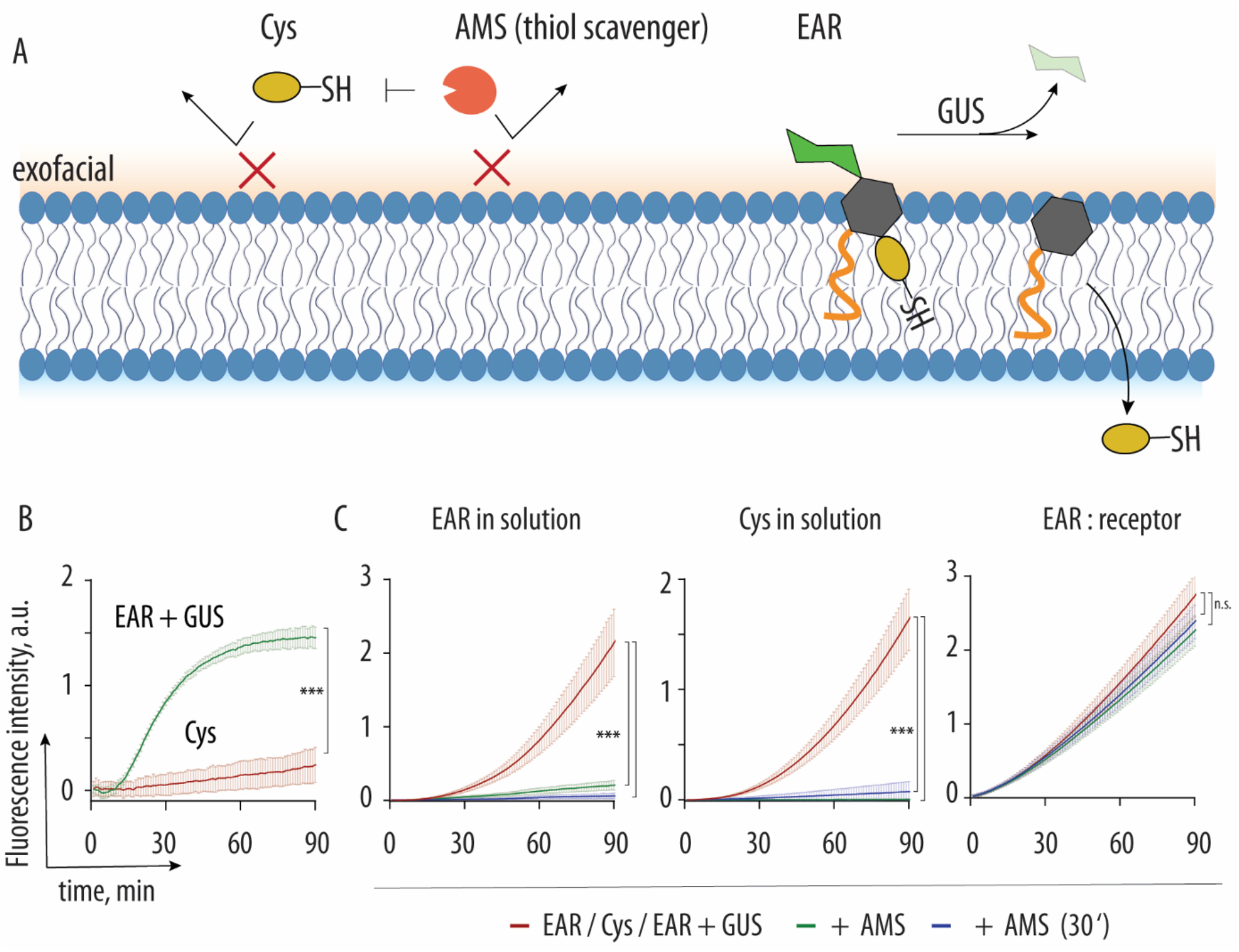
(A) Schematic illustration of transmembrane signalling by EAR or by soluble L-cysteine, in the presence of an exofacial thiol trap molecule, AMS; (B) experimental data for transmembrane activation of the papain zymogen using soluble L-cysteine or GUS-activated EAR, taken at matched concentration; (C) thiol deactivation activity of AMS on L-cysteine or EAR in solution or as a bilayer anchored EAR performing as a bionic receptor. In panels B and C, presented data is based on at least three independent experiments and presented as mean ± SD. In panel B the statistical significance was established via an unpaired t-test and in panel C statistical significance was obtained via a one-way ANOVA test with multiple comparison using the software Graphpad Prism, *** *p* < 0.001; n.s. = non significant.

Finally, we also aimed to gain a better insight into the EAR anchoring within the lipid bilayer and specifically to probe the position of the conjugated cysteine. For this, we used a lipid bilayer-impermeable maleimide (4-acetamido-4-maleimidylstilbene-2,2-disulfonic acid, AMS) which is routinely used in the “cysteine accessibility mapping” as a probe to selectively trap extracellular (exofacial) thiols.^24^ To avoid spectral overlap, enzymatic activity readout was conducted using fluorescein diacetate (typically used as a substrate for esterases but which has proven in our hands to be within the substrate scope of papain). In solution, both EAR and L-cysteine readily activated papain zymogen, whereas addition of AMS was immediately followed by a near-complete inhibition of thiol-disulfide exchange and suppression of zymogen activation to negligible levels (Figure 4C). In stark contrast, within the lipid bilayer, the EAR molecule revealed only minor inhibition by AMS, which illustrates that the EAR thiol is not accessible to the AMS thiol inhibition treatment. This observation strongly suggests, that the EAR cysteine thiol is localized within the lipid bilayer. EAR deactivation was insignificant even when AMS was pre-incubated with the EAR-containing liposomes for 30 minutes prior to the addition of GUS, illustrating the conjugated cysteine is fast anchored within the bilayer.

Taken together, results of this study present the design and performance of a chemical, bionic signalling receptor that responds to a biological primary messenger, generates a biological secondary messenger, and affords a triggered biological event, namely activation of a zymogen for downstream signalling. Similarity of this bionic receptor with biological and chimeric counterparts is in having a well-defined exofacial part and in an in-membrane signal transducer, in responding to a biological primary messenger, and in activation of an enzyme for downstream signalling. For signal transduction, we use a self-immolative molecule, which affords a triggered, traceless release of a secondary messenger for activation of the encapsulated zymogen. The receptor molecule released the secondary messenger directly into the host vesicle and not into the solution bulk, which is the key attribute to postulate receptor-like mechanism of signal transduction. We believe that our work goes well beyond the state of art in that chemical transmembrane signalling performed in this work is activated by a biological input, and affords a biologically relevant output, which has not been achieved previously. We anticipate that these results will significantly empower the fields of artificial and synthetic cell engineering. ^25–28^

## Supporting information

Experimental section and Supporting Information

## Acknowledgements

We wish to acknowledge funding from the Lundbeck Foundation (Grant No R164-2013-15291), the Independent Research Fund Denmark (DFF FNU Grant No 0135-00162B), and the Novo Nordisk Foundation (Grant No NNF20OC0062131).

