## Supplementary material for "Transmembrane signalling by a bionic receptor: biological input and output, chemical mechanism of signal transduction": Experimental section and Supporting Information

### 1. Chemistry section

#### 1.1 General information

All chemicals and solvents were acquired from Sigma Aldrich and used without further purification unless otherwise stated. *N,N*-Dimethylformamide (DMF), trimethylamine (TEA), and methanol were obtained in anhydrous state. The solvents acetonitrile, dichloromethane, and tetrahydrofurane (THF) were dried over aluminum oxide using a MBraun SP800 purification system. The deuterated solvents used for $^{1}$H-NMR and $^{13}$C-NMR analysis were purchased from EurisoTop. Thin layer chromatography (TLC) analysis was performed using silica coated aluminum foil plates (Merck Kieselgel 60 F254) and the plates were analyzed by visualizing either by UV irradiation and/or staining with $KMnO_4$. High purity grade silica gel (w/Ca, ~0.1%, 230-400 mesh particle size, 60Å pore size) was the stationary phase, which was used for silica column chromatography.

A Bruker BioSpin Gmbh 400 MHz spectrometer or a Varian Mercury 400 MHz spectrometer were used for recording nuclear magnetic resonance (NMR) spectra as either $^{1}$H-NMR 400 MHz or $^{13}$C-NMR 101 MHz. The spectra were referenced to the solvent peak.

High-resolution mass spectrometry (HR-MS) was performed using a Bruker Maxis Impact-TOF-MS with electrospray ionization (ESI) and analyzed with Bruker DataAnalysis.

High-performance liquid chromatography (HPLC) experiments were conducted with an Agilent 1260 Infinity II connected to an EC-C18 column with particle size of 2.7 µm, length of 100 mm and diameter of 4.6 mm. The mobile phase was a combination of ultrapure water with trifluoroacetic acid (TFA, 0.1% v/v%, eluent A) and HPLC grade acetonitrile with TFA (0.1% v/v%, eluent B). HPLC experiments were performed with following method; started at 5% eluent B content and the eluent B content was gradually increased to 100% within 18 minutes and kept at 100% content for additionally 13 minutes (total time of 31 minutes). UV was measured with the wavelengths $\lambda$ = 210 nm and $\lambda$ = 254 nm.

#### 1.2 Synthesis overview

Below is the full overview of the 11-step synthesis to isolate the EAR (compound **12**) starting from *O*-per-acetylated methyl glucuronate (compound **1**).

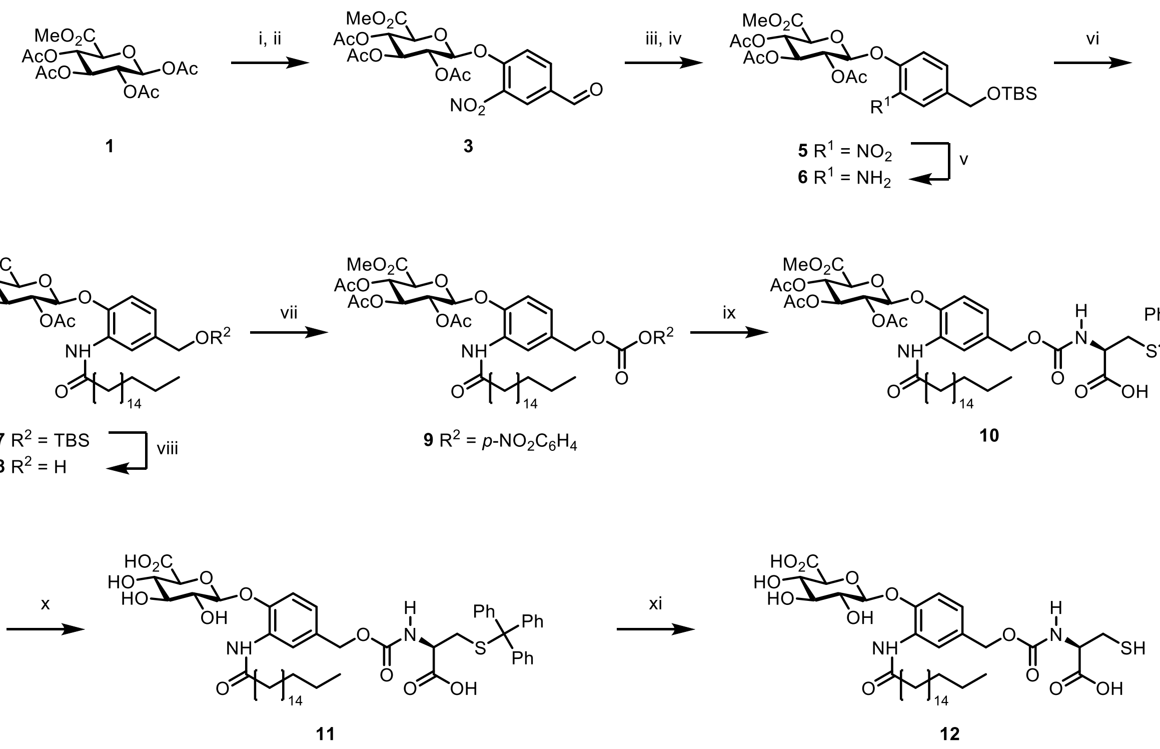

**Figure S1**: Overview of the synthesis of compound **12**. Conditions and reagents; i) HBr/AcOH, $CH_2Cl_2$, 0ºC to r.t., 4 h, 94%, ii) 4-hydroxy-3-nitrobenzaldehyde, $Ag_2O$, powdered 3Å molecular sieves, $CH_3CN$, r.t., 22 h, 68%, iii) $NaBH_4$, silica gel, isopropanol/$CHCl_3$, 0ºC, 1.5 h, 77%, iv) TBSCl, DMAP, imidazole, DMF, r.t, 41.5 h, 82%, v) ammonium formate, Pd/C, abs. EtOH, r.t., 2.5 h, 84%, vi) stearoyl chloride, TEA, $CH_2Cl_2$, 0ºC to r.t., 4 .5h, 96%, vii) TEA·3HF, THF, 0ºC to r.t., 42 h, 59%, viii) 4-nitrophenyl chloroformate, TEA, $CH_2Cl_2$, 0ºC to r.t., 20 h, 75%, ix) *S*-Trityl-L-cysteine, TEA, $CH_2Cl_2$, 0ºC to r.t., 3 h, 80%, x) NaOMe, NaOH, MeOH, 0ºC to r.t., 34%, xi) TFA, (*i*-Pr)$_3$SiH, $CH_2Cl_2$, 0ºC to r.t., 2 h, 16%.

#### 1.3 Synthetic protocols

##### 1.3.1 Synthesis of Compound 2

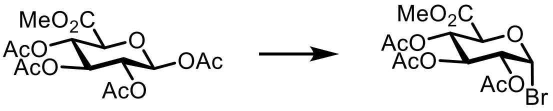

A solution of Compound **1** (2.00 g, 1 equiv.) in $CH_2Cl_2$ (10 mL) was prepared in a round-bottomed flask with a stirring bar under argon atmosphere. The solution was cooled to 0°C and HBr/AcOH (20 mL) was added dropwise under stirring. The resulting brown mixture was allowed to heat to room temperature and stirred for 4 hours until full conversion was observed on TLC. The crude reaction mixture was quenched by pouring onto ice followed by washing of the organic phase with ice water (3x30 mL) and saturated $NaHCO_3$ (3x30 mL). The $CH_2Cl_2$ layer was dried over $MgSO_4$, filtered, reduced on a rotary evaporator, and dried under vacuum to yield the desired compound **2** as a brown syrup (1.99 g, 94%). The compound was used right away for the next step due to instability.

**HR-MS** (ESI): calcd. for $[C_{13}H_{17}BrO_9 + H]^+$: 397.0129, found 397.0713.

**$^1$H-NMR** (400 MHz, Chloroform-*d*) δ 6.64 (d, $J$ = 4.0 Hz, 1H), 5.61 (t, $J$ = 9.7 Hz, 1H), 5.27 – 5.20 (m, 1H), 4.85 (dd, $J$ = 10.0, 4.1 Hz, 1H), 4.57 (d, $J$ = 10.3 Hz, 1H), 3.76 (s, 3H), 2.10 (s, 3H), 2.05 (s, 3H), 2.05 (s, 3H).

The $^1$H-NMR and HR-MS matched the previously published spectrum (Monge et al.; *Adv Sci.* **2021**, DOI: 10.1002/advs.202004432) and this product was used in the next step without additional characterization.

##### 1.3.2 Synthesis of Compound 3

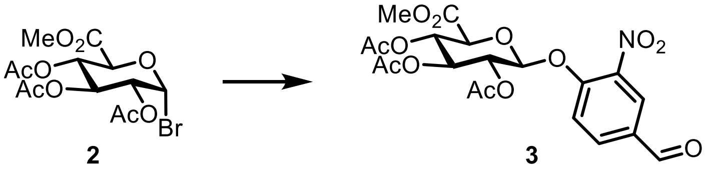

A Schlenk flask with a stirring bar was flame-dried under an atmosphere of argon using Schlenk technique. A suspension of compound **2** (1.99 g, 1 equiv.), 4-hydroxy-3-nitrobenzaldehyde (1.54, 2 equiv.), and powdered 3Å molecular sieves (1.1 g) in $CH_3CN$ (55 mL) was prepared and left stirring. After 30 minutes, $Ag_2O$ (2.14 g, 2 equiv.) was further added to the Schlenk flask and the reaction mixture was left stirring at room temperature in the dark until full conversion was observed on TLC (22 hours). The crude mixture was filtered over a plug of Celite® followed by washing with $CH_2Cl_2$. The filtrate was isolated and the volume was reduced by rotary evaporation. The resulting crude syrup was purified by silica column chromatography eluting with pentane:EtOAc 3:1 to 3:2 yielding the wanted compound **3** (1.65 g, 68%).

**HR-MS** (ESI): calcd. for $[C_{20}H_{21}NO_{13}+Na]^+$: 506.0905, found 506.0921, calcd. for $[C_{20}H_{21}NO_{13}+K]^+$: 522.0645, found 522.0637.

**$^1$H-NMR** (400 MHz, Acetone-$d_6$) δ 10.05 (s, 1H), 8.40 (d, $J$ = 2.0 Hz, 1H), 8.21 (dd, $J$ = 8.7, 2.0 Hz, 1H), 7.78 (d, $J$ = 8.7 Hz, 1H), 5.87 (d, $J$ = 7.4 Hz, 1H), 5.46 (t, $J$ = 9.2 Hz, 1H), 5.35 – 5.22 (m, 2H), 4.73 (d, $J$ = 9.4 Hz, 1H), 3.68 (s, 3H), 2.04 (s, 6H), 2.02 (s, 3H), 2.00 (s, 3H).

**$^{13}$C-NMR** (101 MHz, Acetone) δ 190.4, 170.3, 170.1, 169.5, 167.7, 153.8, 142.1, 135.3, 132.7, 126.9, 119.1, 99.3, 73.1, 71.8, 71.0, 69.7, 53.2, 20.6, 20.6, 20.6.

##### 1.3.3 Synthesis of Compound 4

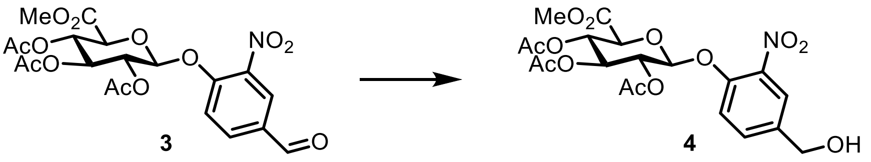

Compound **3** (1.50 g, 1 equiv.) was dissolved in a mixture of $CHCl_3$ (26 mL) and isopropanol (6.5 mL) in a flame-dried round-bottomed flask under an atmosphere of argon. The solution was cooled to 0°C and silica gel (1.37 g) was further added to the mixture. After 10 minutes of stirring, $NaBH_4$ (0.235 g, 2 equiv) was added in one portion to the reaction mixture and left stirring for 1.5 hours. The resulting crude was filtered over a plug of Celite® and washed with $CH_2Cl_2$. The filtrate was collected and washed with brine (3 x 20 mL) followed by drying over $MgSO_4$, filtration and concentration under reduced pressure. The resulting white solid was dried under vacuum to yield the desired compound **4** without further purification (1.16 g, 77%)

**HR-MS** (ESI): calcd. for $[C_{20}H_{23}NO_{13} + Na]^+$ 508.1061, found 508.1076. calcd. for $[C_{20}H_{23}NO_{13} + K]^+$ 524.0801, found 524.0808.

**$^1$H-NMR** (400 MHz, Chloroform-$d$) δ 7.81 (d, $J$ = 2.1 Hz, 1H), 7.53 (dd, $J$ = 8.6, 2.1 Hz, 1H), 7.36 (d, $J$ = 8.6 Hz, 1H), 5.39 – 5.22 (m, 3H), 5.19 (d, $J$ = 6.9 Hz, 1H), 4.72 (d, $J$ = 5.7 Hz, 2H), 4.20 (d, $J$ = 9.0 Hz, 1H), 3.74 (s, 3H), 2.12 (s, 3H), 2.06 (s, 3H), 2.05 (s, 3H).

The 1H-NMR and HR-MS matched the previously published spectrum (Monge et al.; Adv Sci. 2021, DOI: 10.1002/advs.202004432) and this product was used in the next step without additional characterization.

##### 1.3.4 Synthesis of Compound 5

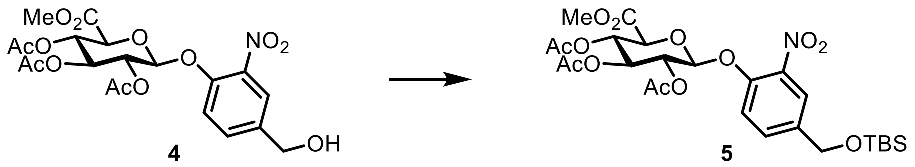

A solution was prepared in a flame-dried flask under an atmosphere of argon by dissolving compound **4** (1.14 g, 1 equiv.) in *N,N*-dimethylformamide (DMF) (11.3 mL). Imidazole (0.956 g, 6 equiv.) and 4-dimethylaminopyridine (DMAP) (0.0715 g, 0.25 equiv.) were further added and the reaction mixture was stirred for 5 minutes. Finally, TBSCl (2.12 g, 6 equiv.) in DMF (8.0 mL) was added dropwise and the reaction mixture was stirred at room temperature until full conversion was observed on TLC (41.5 hours). Upon completion, the crude mixture was diluted with $CH_2Cl_2$ and the organic layer was washed with $NH_4Cl$ (3x50 mL) and brine (2x50 mL), dried over $Na_2SO_4$, filtered, and concentrated *in vacuo*. The purification was performed by silica column chromatography eluting with pentane:diethyl ether 60:40 to 40:60 yielding the wanted compound **5** (1.15 g, 82%).

**HR-MS** (ESI): calcd. for $[C_{26}H_{37}NO_{13}Si + Na]^+$ 622.1926, found 622.1964. calcd. for $[C_{26}H_{37}NO_{13}Si + K]^+$ 638.1666, found 638.1667.

**$^1$H-NMR** (400 MHz, Chloroform-*d*) δ 7.75 (d, *J* = 2.0 Hz, 1H), 7.48 (dd, *J* = 8.6, 2.1 Hz, 1H), 7.34 (d, *J* = 8.6 Hz, 1H), 5.33 (ddt, *J* = 11.3, 6.7, 3.6 Hz, 3H), 5.17 (d, *J* = 6.9 Hz, 1H), 4.72 (s, 2H), 4.22 – 4.12 (m, 1H), 3.75 (s, 3H), 2.13 (s, 3H), 2.06 (s, 3H), 2.05 (s, 3H), 0.94 (s, 9H), 0.11 (s, 6H).

The $^1$H-NMR and HR-MS matched the previously published spectrum (Monge et al.; *Adv Sci.* **2021**, DOI: 10.1002/advs.202004432) and this product was used in the next step without additional characterization.

##### 1.3.5 Synthesis of Compound 6

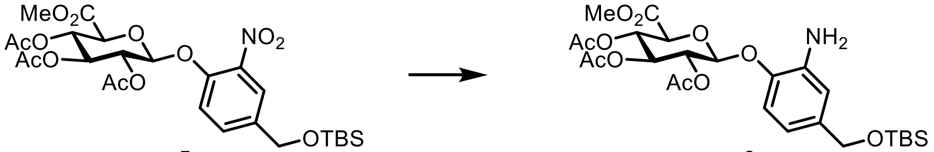

Compound **5** (0.380 g, 1 equiv.) was added to a round-bottom flask with a stirring bar and suspended in absolute ethanol (30 mL). The suspension was purged with argon for 10 minutes and kept under an atmosphere of argon. In one portion, Pd/C (0.053 g, 0.7 equiv.) and ammonium formate (0.160 g, 4 equiv.) was added to the reaction mixture and the resulting suspension was stirred at room temperature for 2.5 hours. Upon completion of the reaction, the mixture was filtered through a plug of Celite® and the volume of filtrate was reduced *in vacuo*. The crude was diluted with EtOAc and washed with brine (3 x 20 mL), dried over $Na_2SO_4$ and filtered. The filtrate was concentrated under reduced pressure to yield the wanted compound **6** (0.305 g, 84%).

**HR-MS** (ESI): $[C_{26}H_{39}NO_{11}Si+H]^+$ calcd. 570.2365, found 570.2404. $[C_{26}H_{39}NO_{11}Si+Na]^+$ calcd. 592.2184, found 592.2204.

**$^1$H-NMR** (400 MHz, Chloroform-*d*) δ 6.87 (d, *J* = 8.2 Hz, 1H), 6.68 (d, *J* = 1.9 Hz, 1H), 6.61 (dd, *J* = 8.2, 2.0 Hz, 1H), 5.31 (qd, *J* = 10.0, 9.3, 3.3 Hz, 3H), 5.00 (d, *J* = 7.1 Hz, 1H), 4.60 (s, 2H), 4.14 (d, *J* = 8.8 Hz, 1H), 3.75 (s, 3H), 2.08 (s, 3H), 2.05 (s, 3H), 2.04 (s, 3H), 0.93 (s, 9H), 0.08 (s, 6H), residual EtOAc (4.12, 2.05, 1.26).

The $^1$H-NMR and HR-MS matched the previously published spectrum (Monge et al.; *Adv Sci.* **2021**, DOI: 10.1002/advs.202004432) and this product was used in the next step without additional characterization.

##### 1.3.6 Synthesis of Compound 7

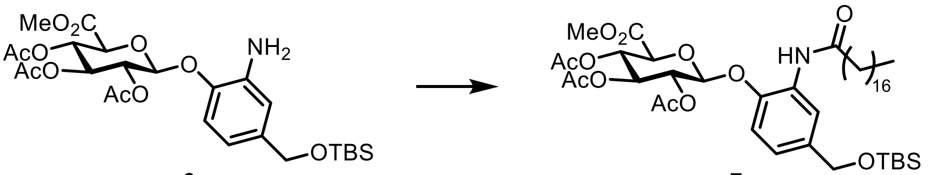

Compound **6** (0.305 g, 1 equiv.), triethyl amine (TEA) (0.149 mL, 2 equiv.) and dry $CH_2Cl_2$ (10 mL) were added to a flame-dried flask with a stirring bar under an atmosphere of argon. The mixture was cooled to 0°C in an ice bath followed by dropwise addition of stearoyl chloride (0.211 g, 1.1 equiv.) in dry $CH_2Cl_2$ (4 mL). The reaction was left stirring at room temperature for 4.5 hours until full conversion was observed on TLC. Upon completion, water (12 mL) was added to the reaction followed by extraction with $CH_2Cl_2$ (3 x20 mL). The organic layers were dried over $MgSO_4$, filtered and the solvent removed under reduced pressure. The resulting crude was purified by silica column chromatography by eluting with pentane:EtOAc 9:1 to 8:2 to yield the pure compound **7** (0.433 g, 96%).

**HR-MS** (ESI): $[C_{44}H_{73}NO_{12}Si+Na]^+$ calcd. 858.4794, found 858.4815.

**$^1$H-NMR** (400 MHz, Chloroform-*d*) δ 8.33 (s, 1H), 7.86 (s, 1H), 7.05 (dd, *J* = 8.4, 2.0 Hz, 1H), 6.90 (d, *J* = 8.4 Hz, 1H), 5.47 – 5.20 (m, 3H), 5.04 (d, *J* = 7.5 Hz, 1H), 4.67 (s, 2H), 4.17 (d, *J* = 9.6 Hz, 1H), 3.75 (s, 3H), 2.40 (td, *J* = 7.4, 3.0 Hz, 2H), 2.08 (s, 3H), 2.07 (s, 3H), 2.05 (s, 3H), 1.70 (q, *J* = 7.3 Hz, 2H), 1.25 (s, 28H), 0.93 (s, 9H), 0.87 (t, *J* = 6.8 Hz, 3H), 0.09 (s, 6H), residual EtOAc (4.12, 2.05, 1.26).

The $^1$H-NMR and HR-MS matched the previously published spectrum (Monge et al.; *Adv Sci.* **2021**, DOI: 10.1002/advs.202004432) and this product was used in the next step without additional characterization.

##### 1.3.7 Synthesis of Compound 8

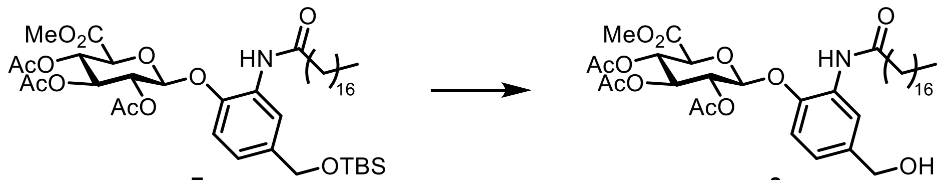

Compound **7** (0.433 g, 1 equiv.) and tetrahydrofurane (THF) (4.8 mL) were mixed in and flame-dried flask under and atmosphere of argon and cooled to 0°C. TEA·3HF (0.110 mL, 1.3 equiv.) in THF (1.7 mL) was slowly added and the reaction mixture was left at room temperature for 42 hours. $CH_2Cl_2$ (30 mL) was added to the reaction mixture and washed with saturated $NH_4Cl$ (2x20 mL) and brine (2x20 mL). The organic layer was dried over $MgSO_4$, filtered and the solvent was removed under reduced pressure. The purification was accomplished by silica column chromatography eluting with pentane:EtOAc 6:4 to 5:5, which afforded the desired compound **8** (0.219 g, 59%).

**HR-MS** (ESI): $[C_{38}H_{59}NO_{12}+H]^+$ calcd. 722.4110, found 722.4121. $[C_{38}H_{59}NO_{12}+Na]^+$ calcd. 744.3929, found 744.3923.

**$^1$H-NMR** (400 MHz, Chloroform-*d*) δ 8.43 (d, *J* = 1.8 Hz, 1H), 7.89 (s, 1H), 7.06 (dd, *J* = 8.3, 2.0 Hz, 1H), 6.92 (d, *J* = 8.3 Hz, 1H), 5.48 – 5.22 (m, 3H), 5.06 (d, *J* = 7.5 Hz, 1H), 4.63 (d, *J* = 6.0 Hz, 2H), 4.19 (d, *J* = 9.6 Hz, 1H), 3.76 (s, 3H), 2.42 (td, *J* = 7.4, 3.2 Hz, 2H), 2.08 (s, 3H), 2.07 (s, 3H), 2.06 (s, 3H), 1.72 (p, *J* = 7.6, 7.1 Hz, 2H), 1.25 (s, 28H), 0.88 (t, *J* = 6.8 Hz, 3H).

##### 1.3.8 Synthesis of Compound 9

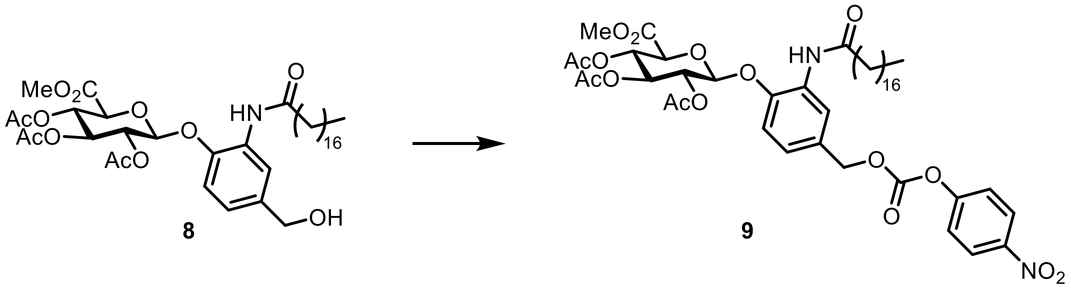

A solution was prepared in a flame-dried flask of compound **8** (0.100 g, 1 equiv.) and TEA (0.058 mL, 3 equiv.) in dry $CH_2Cl_2$ (0.7 mL) under argon atmosphere. The mixture was cooled to 0°C followed by dropwise addition of 4-nitrophenyl chloroformate (0.039 g, 1.4 equiv.) in dry $CH_2Cl_2$ (1 mL). The reaction mixture was stirred at room temperature for 20 hours until the starting material was consumed. The crude was absorbed onto Celite® and subjected to silica column chromatography purification eluting with pentane:EtOAc 8:2 to 7:3 yielding the pure compound **9** (0.092 g, 75%).

**HR-MS** (ESI): $[C_{45}H_{62}N_2O_{16}+Na]^+$ calcd. 909.3991, found 909.3953.

**$^1$H-NMR** (400 MHz, Chloroform-*d*) δ 8.59 (d, *J* = 1.9 Hz, 1H), 8.34 – 8.20 (m, 2H), 7.91 (s, 1H), 7.43 – 7.34 (m, 2H), 7.09 (dd, *J* = 8.4, 2.1 Hz, 1H), 6.95 (d, *J* = 8.4 Hz, 1H), 5.50 – 5.26 (m, 3H), 5.23 (s, 2H), 5.09 (d, *J* = 7.5 Hz, 1H), 4.21 (d, *J* = 9.6 Hz, 1H), 3.76 (s, 3H), 2.44 (td, *J* = 7.4, 4.1 Hz, 2H), 2.09 (s, 3H), 2.08 (s, 3H), 2.06 (s, 3H), 1.73 (p, *J* = 7.6, 7.1 Hz, 2H), 1.25 (s, 28H), 0.88 (t, *J* = 6.8 Hz, 3H).

The $^1$H-NMR and HR-MS matched the previously published spectrum (Monge et al.; *Adv Sci.* **2021**, DOI: 10.1002/advs.202004432) and this product was used in the next step without additional characterization.

##### 1.3.9 Synthesis of compound 10

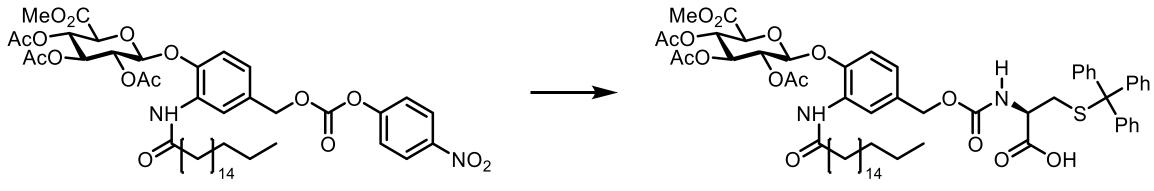

9 10

*S*-trityl-L-cysteine (0.033 g, 1.2 equiv.) and triethylamine (TEA) (0.027 g, 3 equiv.) were mixed in dry $CH_2Cl_2$ ( 0.7 mL) under an atmosphere of argon and cooled to 0°C. To that suspension was added Compound **9** (0.067 g, 1 equiv.) in dry $CH_2Cl_2$ (0.7 mL) and the mixture was allowed to heat to room temperature. After 3 hours, TLC showed full consumption of the starting material. $CH_2Cl_2$ was removed under reduced mixture and the crude product was directly absorbed onto Celite®. The purification was performed using silica column chromatography eluting with 4-10% MeOH in $CH_2Cl_2$ yielding the wanted compound **10** as a white solid (0.067 g, 80%).

**HR-MS** (ESI) positive: $[C_{61}H_{78}N_2O_{15}S+Na]^+$ calcd. 1133.5015, found 1133.5008. $[C_{61}H_{78}N_2O_{15}S+K]^+$ calcd. 1149.4755, found 1149.4798. $[C_{61}H_{78}N_2O_{15}S+2Na-H]^+$ calcd. 1155.4835, found 1155.4830.

**$^1$H-NMR** (400 MHz, Chloroform-*d*) δ (ppm) 8.37 (s, 1H), 7.91 (s, 1H), 7.38 (d, *J* = 7.7 Hz, 6H), 7.28 – 7.15 (m, 9H), 6.92-6.85 (m, 2H), 5.46 – 5.27 (m, 4H), 5.06 (d, *J* = 7.6 Hz, 1H), 4.98 – 4.84 (m, 2H), 4.23 (d, *J* = 9.7 Hz, 1H), 4.15 – 4.13 (m, 1H), 3.73 (s, 3H), 2.63 (d, *J* = 5.6 Hz, 2H), 2.41 (td, *J* = 7.3, 2.8 Hz, 2H), 2.07-2.06 (m, 9H), 1.70 (p, *J* = 7.3 Hz, 2H), 1.25 (s, 28H), 0.88 (t, *J* = 6.7 Hz, 3H).

**$^{13}$C-NMR** (101 MHz, Chloroform-*d*) δ 172.6, 170.3, 169.9, 169.6, 166.7, 156.2, 144.9, 144.5, 132.4, 129.7, 129.1, 128.1, 127.0, 123.3, 120.3, 114.5, 100.0, 72.5, 71.3, 71.2, 69.4, 67.1, 66.6, 53.2, 37.8, 33.8, 32.1, 29.9, 29.8, 29.7, 29.7, 29.5, 25.8, 22.8, 20.9, 20.7, 20.6, 14.3.

**HPLC**: $t_r$ = 28.18 min.

#### 1.3.10 Synthesis of Compound 11

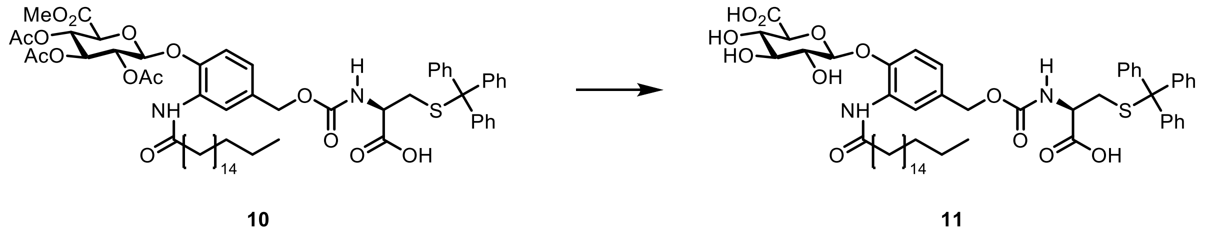

A solution of Compound **10** (0.067 g, 1 equiv.) in dry methanol (1.0 mL) was prepared in a flame-dried flask under an atmosphere of argon. The mixture was cooled to 0°C and NaOMe (0.3 equiv.) was added dropwise. The reaction was allowed to heat to room temperature while it was followed on TLC. After an hour, more NaOMe (0.3 equiv.) was added in the same manner under cooling, since the reaction was still not finished. This was repeated until full conversion was observed on TLC, which happened after three additions (3 hours). Upon completion, the mixture was cooled to 0°C and $H_2O$ (1 mL) and 2 M NaOH (30 µL) were slowly added and the reaction was left for another 10 minutes. The reaction mixture was neutralized with amberlite 120$H^+$ ion exchange resin followed by filtration and washing with methanol. The resulting solution was lyophilized to yield a white powder. The purification was performed by silica column chromatography eluting with 6-10% MeOH and 1% AcOH in $CH_2Cl_2$ followed by MeOH:$CH_3CN$:$H_2O$:EtOAc 1:1:1:7 affording the desired Compound **11** as a white powder (0.020 g, 34%).

**HR-MS** (ESI) positive: $[C_{54}H_{70}N_2O_{12}S+Na]^+$ calcd. 993.4541, found 993.4553. $[C_{54}H_{70}N_2O_{12}S+2Na-H]^+$ calcd. 1015.4361, found 1015.4373.

**HR-MS** (ESI) negative: $[C_{54}H_{70}N_2O_{12}S-H]^-$ calcd. 969.4576, found 969.4458. $[C_{54}H_{70}N_2O_{12}S-H]^-$ calcd. 991.4396, found 991.4278.

**$^1$H-NMR** (400 MHz, DMSO-$d_6$) δ 9.24 (s, 1H), 8.10 (s, 1H), 7.34 – 7.17 (m, 16H), 7.13 (d, $J$ = 8.3 Hz, 1H), 7.03 (d, $J$ = 7.8 Hz, 1H), 6.61 (bs, 1H), 5.75 (bs, 1H), 5.10 (bs, 1H), 4.99 – 4.82 (m, 2H), 4.55 (d, $J$ = 7.1 Hz, 1H), 3.77 (d, $J$ = 4.7 Hz, 1H), 3.29 – 3.21 (m, 2H), 3.20 – 3.12 (m, 1H), 2.46 – 2.34 (m, 3H), 1.84 (s, 2H), 1.60 – 1.51 (m, 2H), 1.23 (s, 28H), 0.88 – 0.81 (m, 3H).

**HPLC**: $t_r$ (**11**) = 24.55 min., $t_r$ (**11** with GUS) = 12.75 min., $t_r$ (trityl cysteine) = 13.05 min.

##### 1.3.11 Synthesis of Compound 12

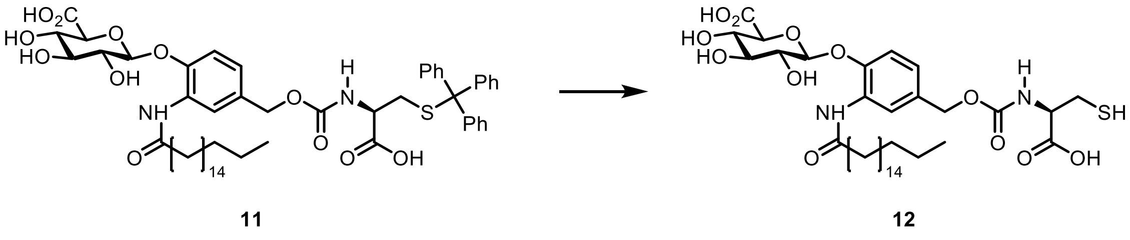

A mixture was prepared of Compound **11** (0.027 g, 1 equiv.) in dry $CH_2Cl_2$ (2.5 mL) under argon atmosphere and the resulting solution was cooled to 0°C. (*i*-Pr)$_3$SiH (0.011 mL, 2 equiv.) and trifluoroacetic acid (TFA) (0.107 mL, 50 equiv.) were added to the reaction and the mixture was allowed to heat to room temperature. The mixture was stirred for 2 hours before full conversion was observed. The volume of the resulting crude was reduced *in vacuo* followed by silica column chromatography purification eluting with 10% MeOH in $CH_2Cl_2$ followed by MeOH:$CH_3CN$:$H_2O$:EtOAc 1:1:1:7. Compound **12** was isolated as a white solid (3.2 mg, 16%).

**HR-MS** (ESI) positive: $[C_{35}H_{56}N_2O_{12}S+Na]^+$ calcd. 751.3446, found 751.3462. $[C_{35}H_{56}N_2O_{12}S+2Na-H]^+$ calcd. 773.3266, found 773.3288.

**HR-MS** (ESI) negative: $[C_{35}H_{56}N_2O_{12}S-H]^-$ calcd. 727.3481, found 727.3458. $[C_{35}H_{56}N_2O_{12}S+Na-2H]^-$ calcd. 749.3301, found 749.3288.

**HPLC**: $t_r$ = 21.58 min.

### 1.4 NMR spectra

#### 1.4.1 NMR spectrum of Compound 2

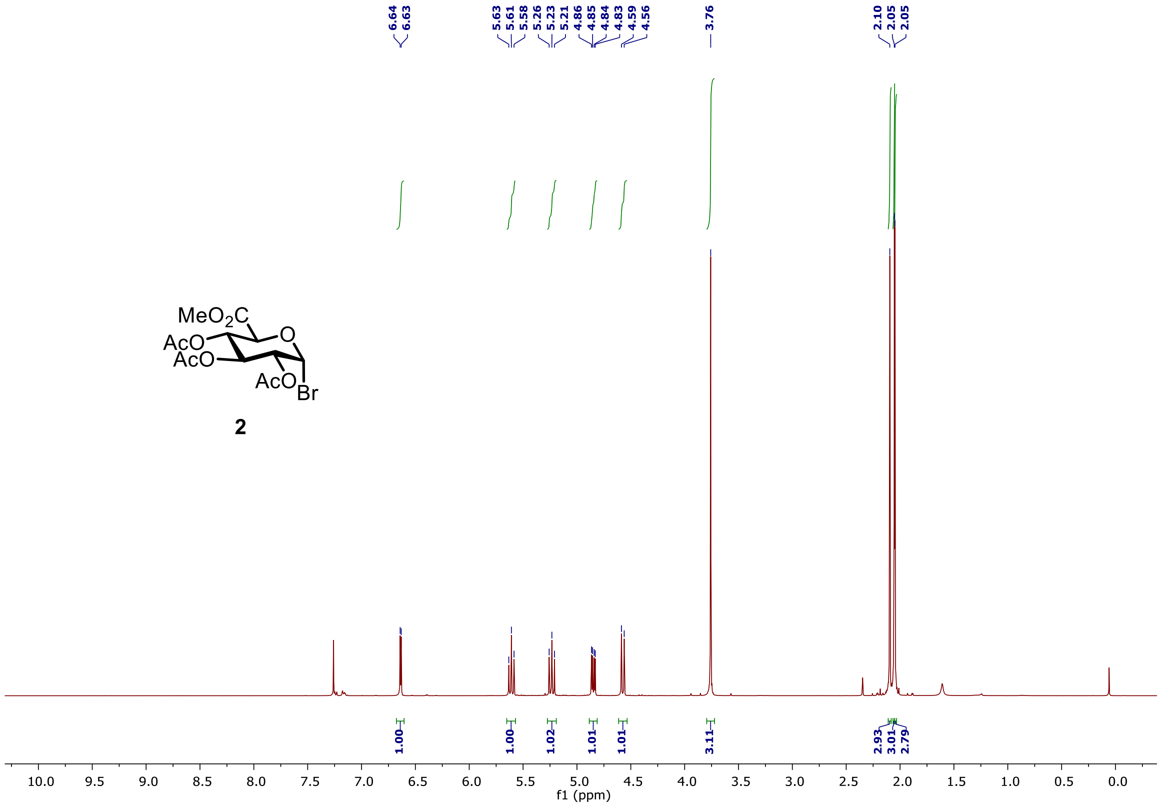

#### 1.4.2 NMR spectra of Compound 3

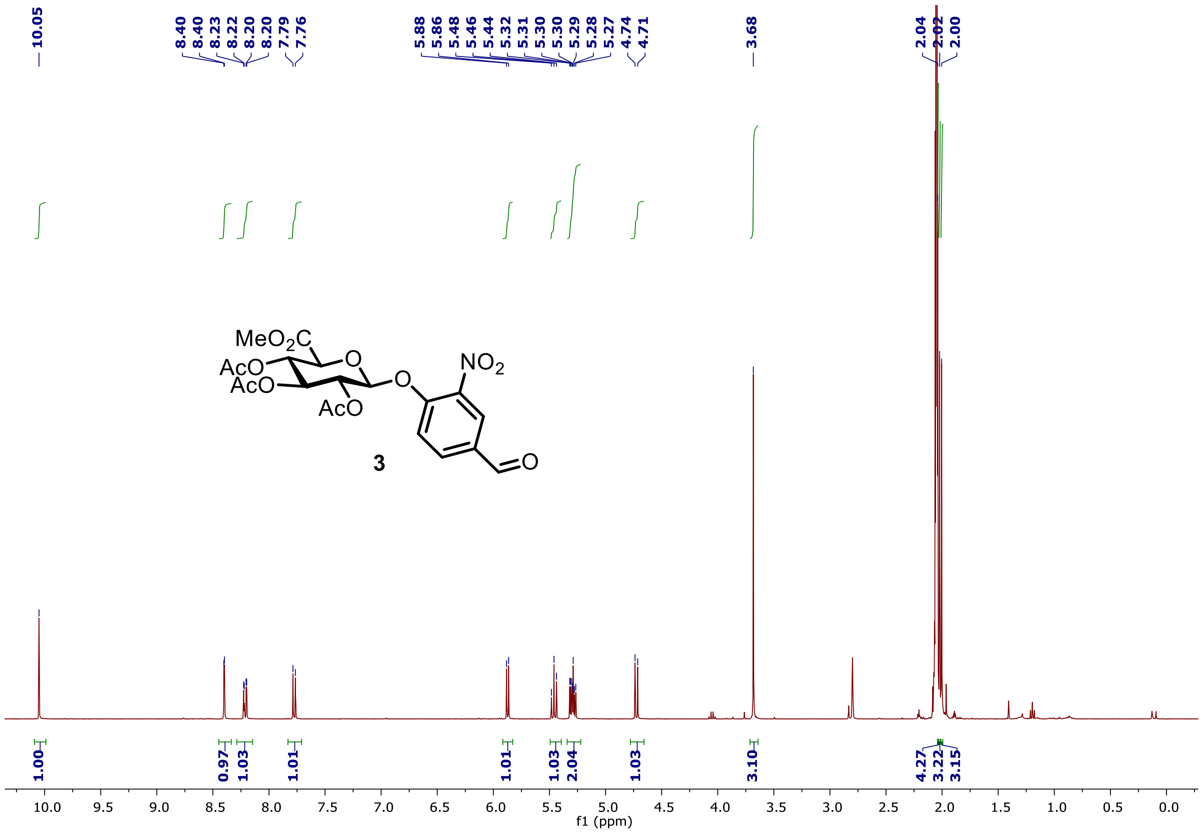

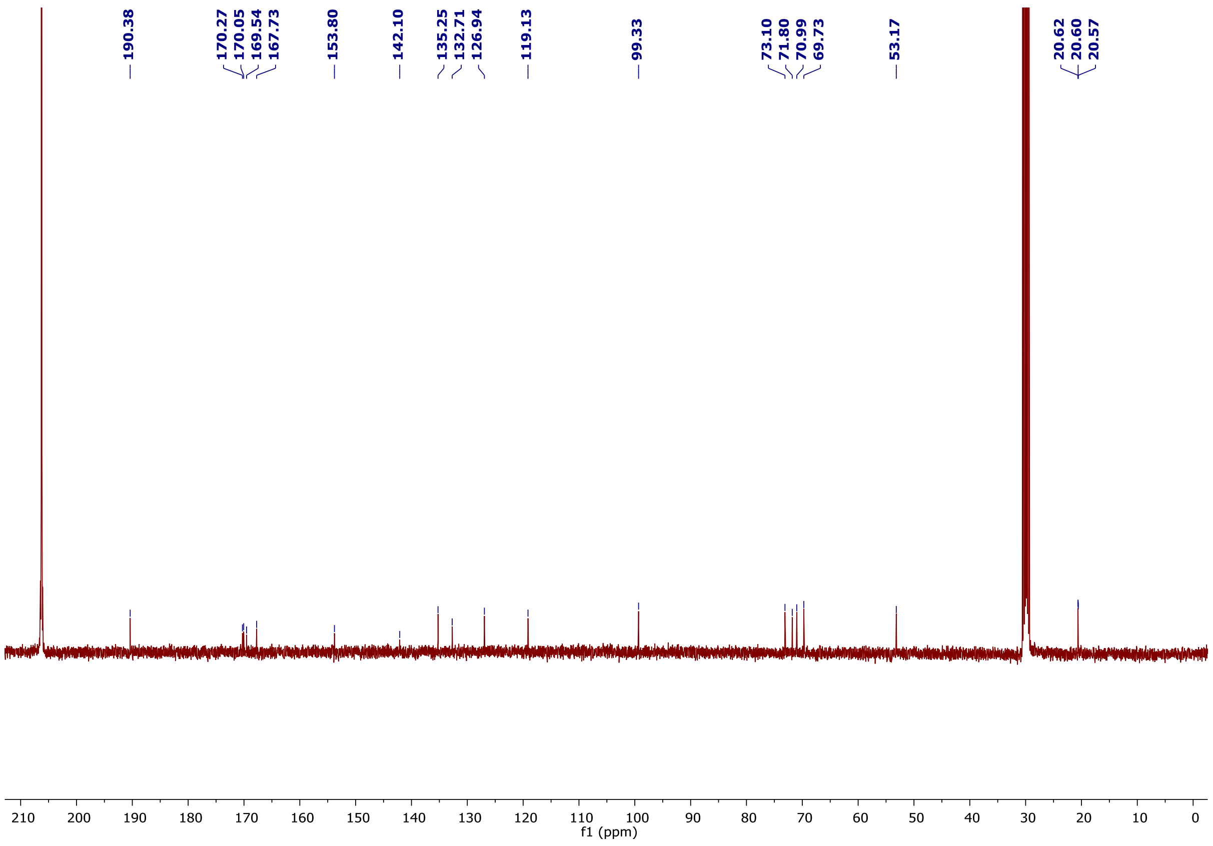

#### 1.4.3 NMR spectrum of Compound 4

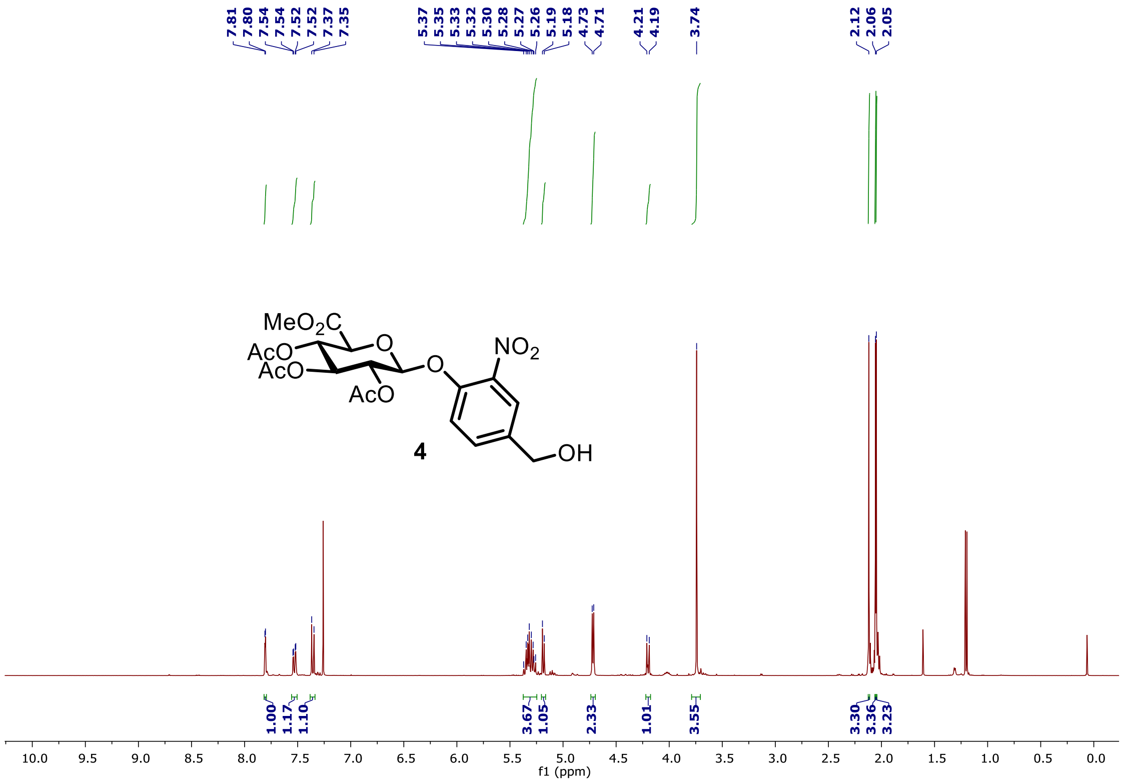

#### 1.4.4 NMR spectrum of Compound 5

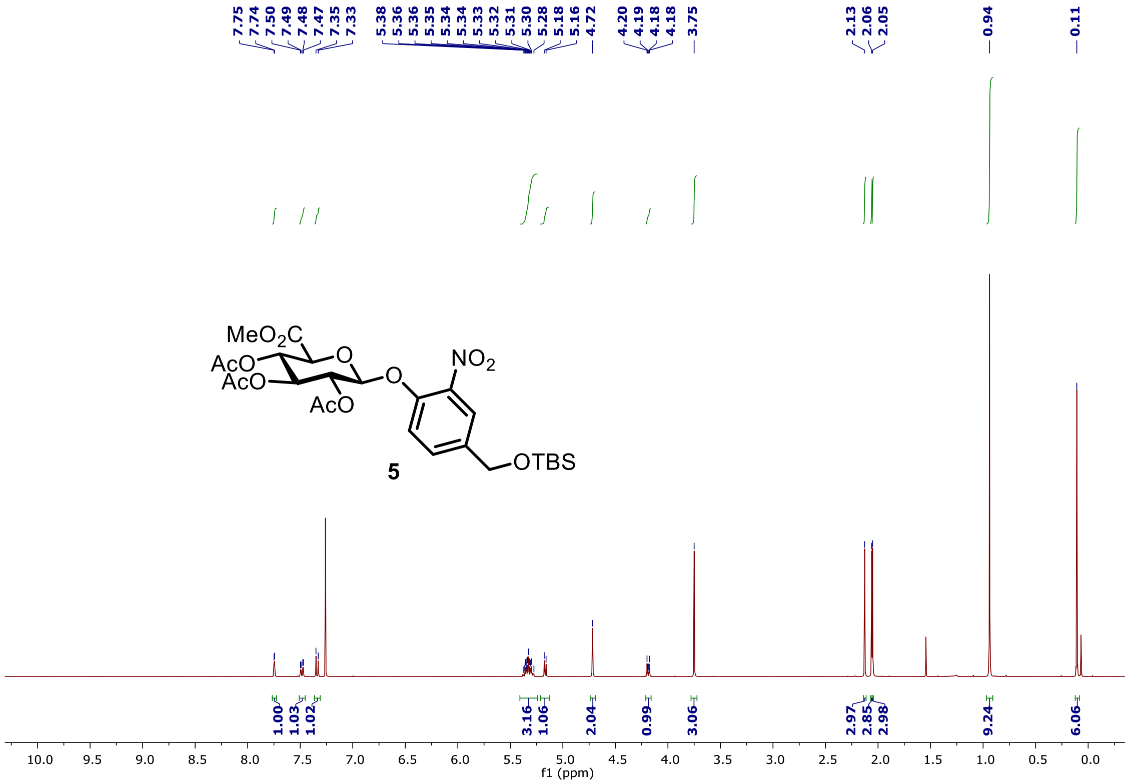

#### 1.4.5 NMR spectrum of Compound 6

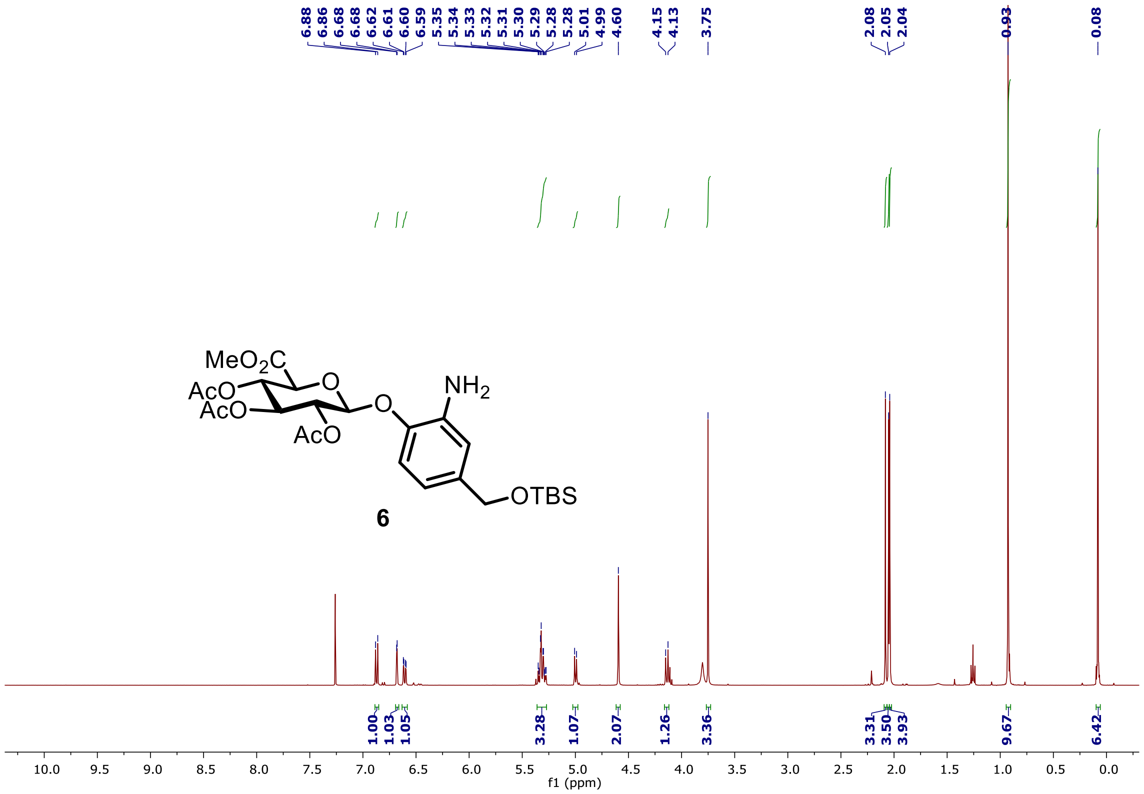

#### 1.4.6 NMR spectrum of Compound 7

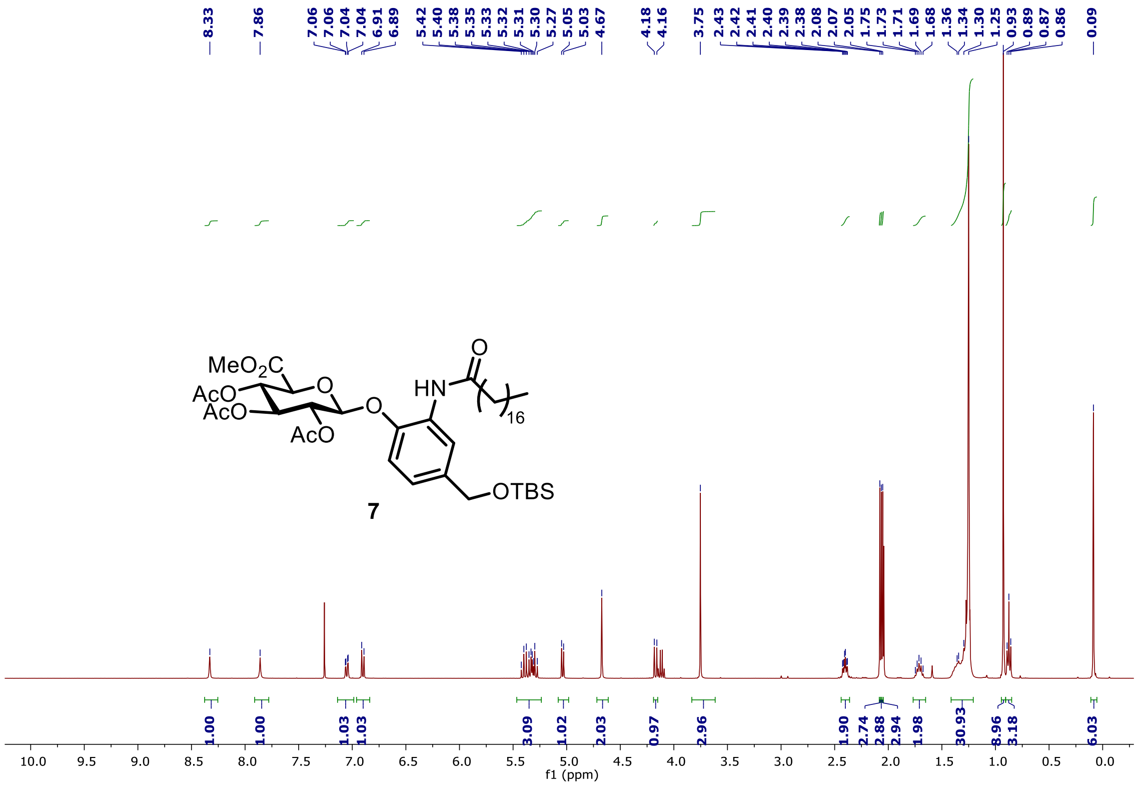

#### 1.4.7 NMR spectrum of Compound 8

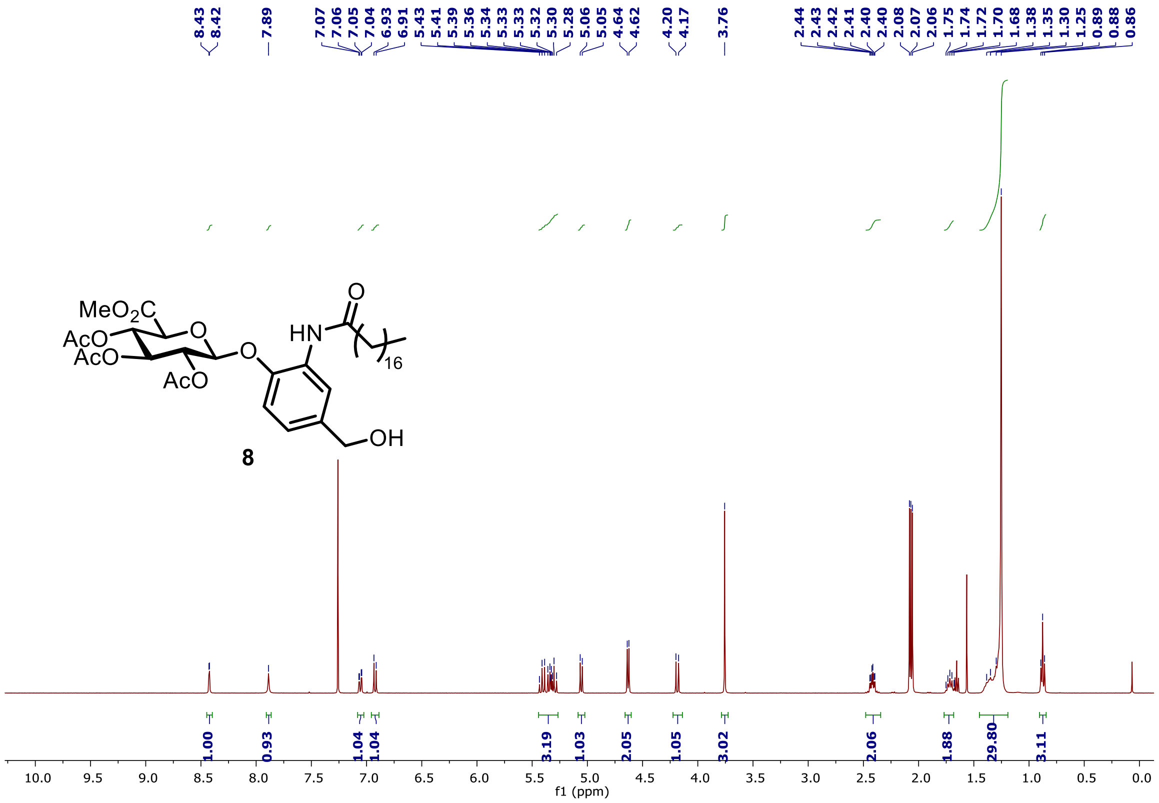

#### 1.4.8 NMR spectrum of Compound 9

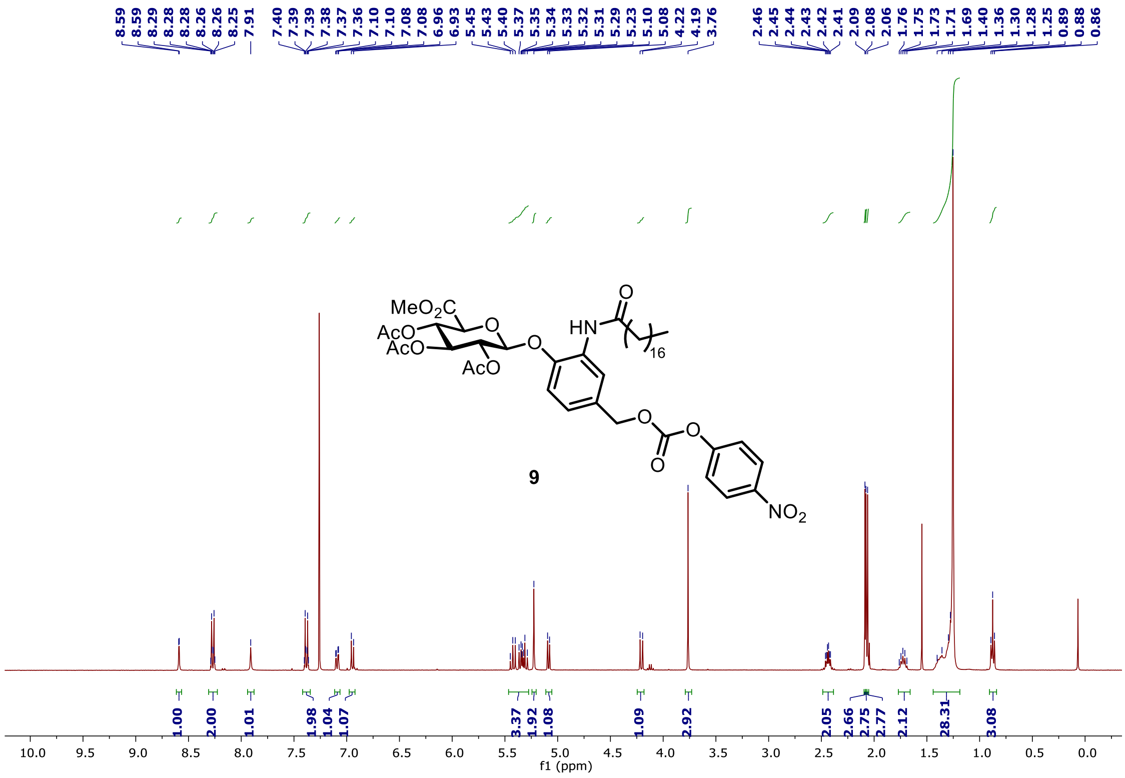

#### 1.4.9 NMR spectra of Compound 10

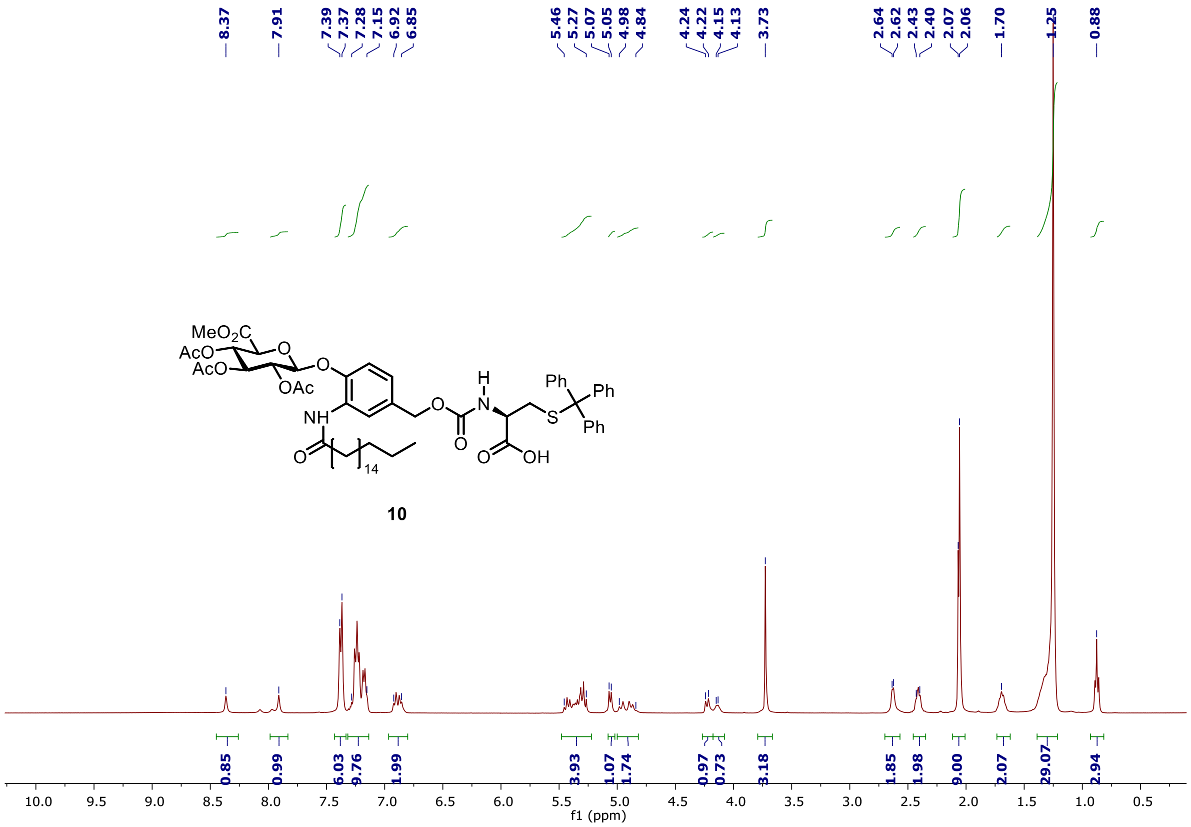

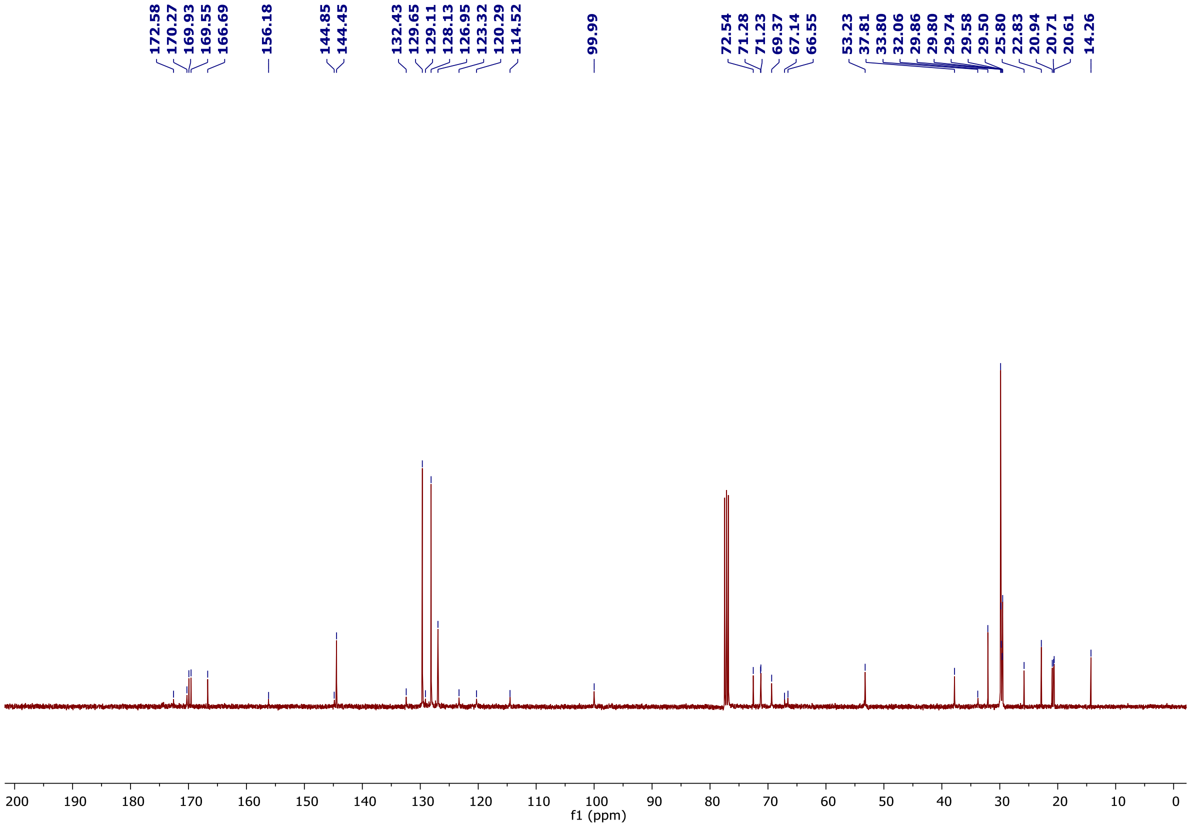

#### 1.4.10 NMR spectrum of Compound 11

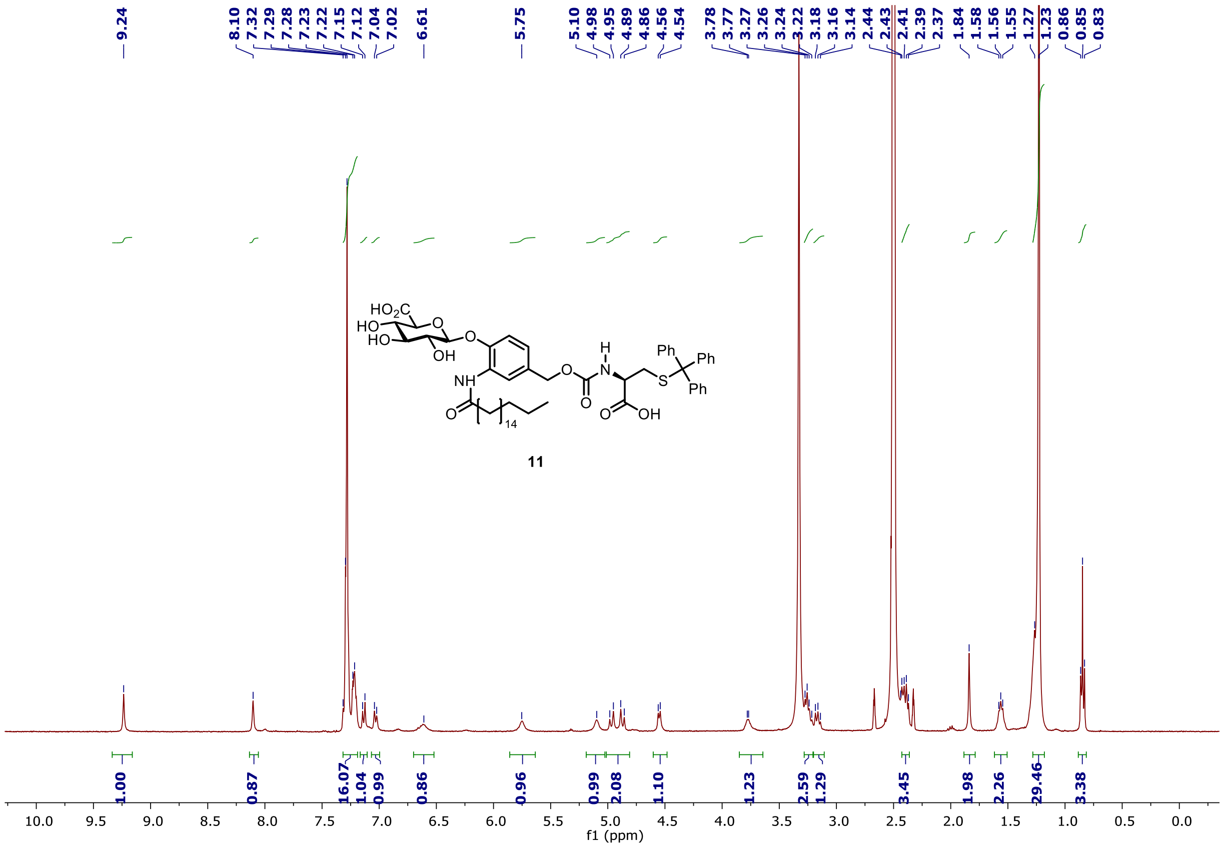

#### 1.5 MS spectra

Below is shown the MS spectra of compound **10**, **11**, and **12**.

##### 1.5.1 MS spectrum of Compound 10

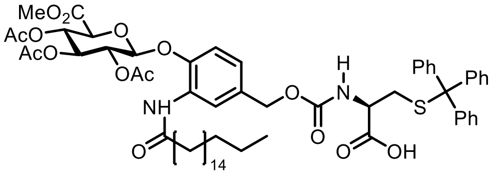

**10**

**Positive mode:**

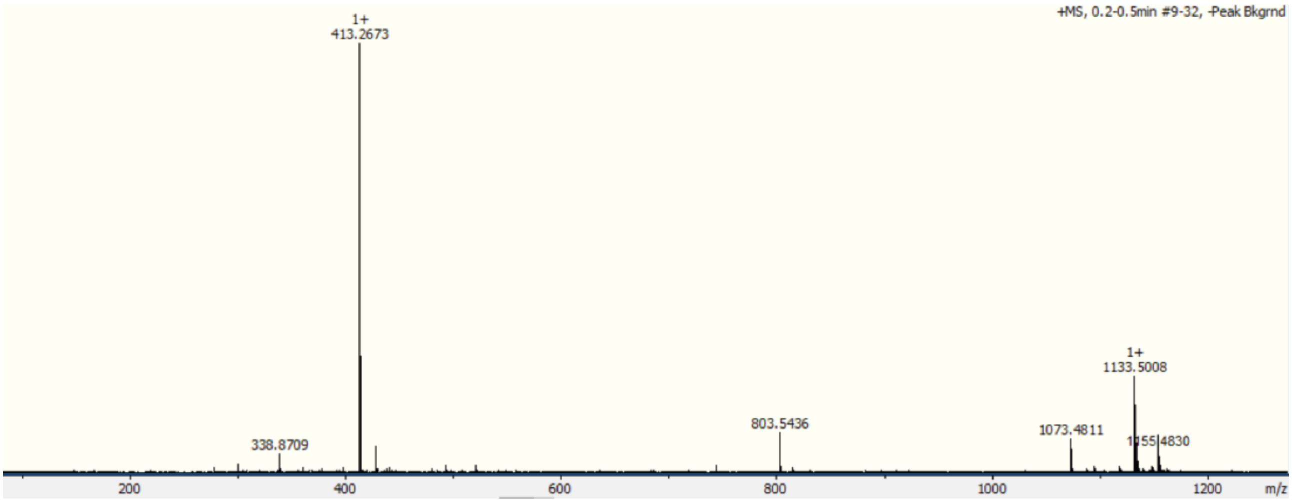

##### 1.5.2 MS spectra of Compound 11

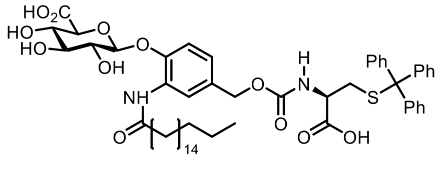

**11**

**Positive mode:**

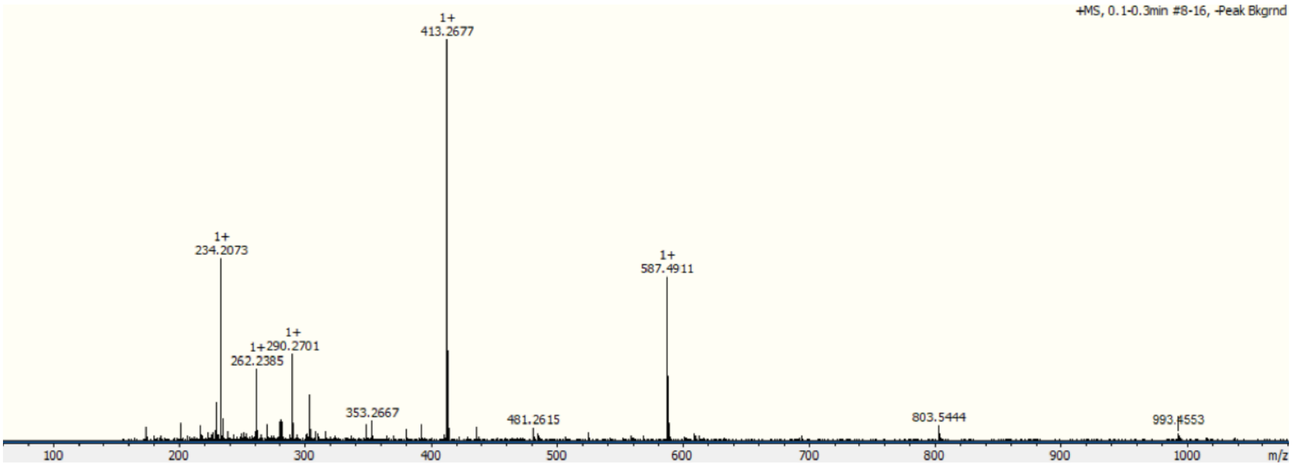

**Negative mode:**

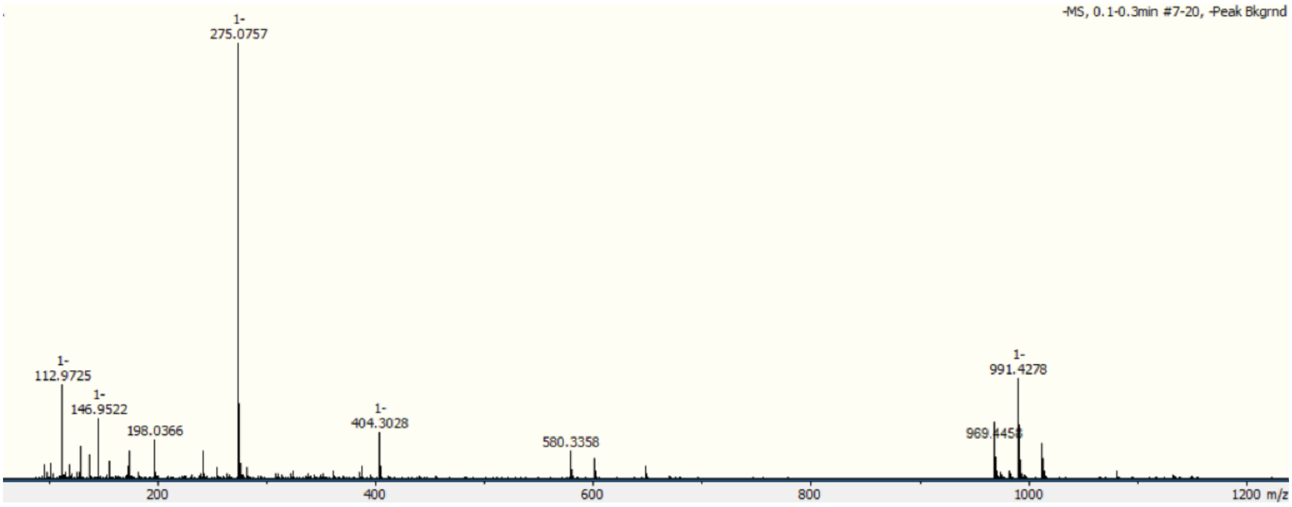

##### 1.5.3 MS spectra of Compound 12

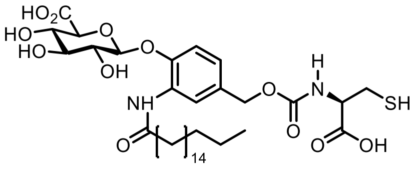

**12**

**Positive mode:**

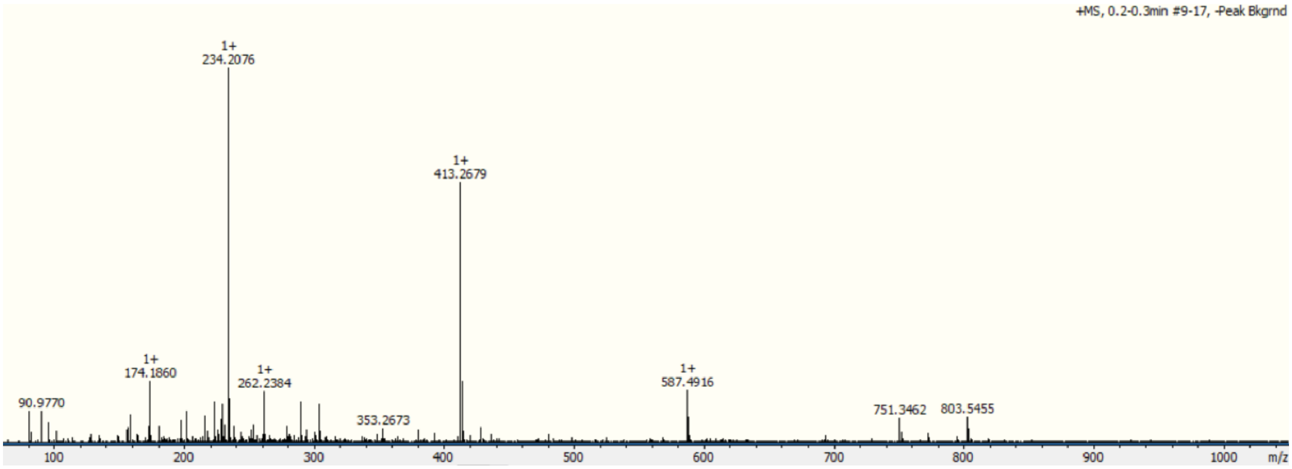

**Negative mode:**

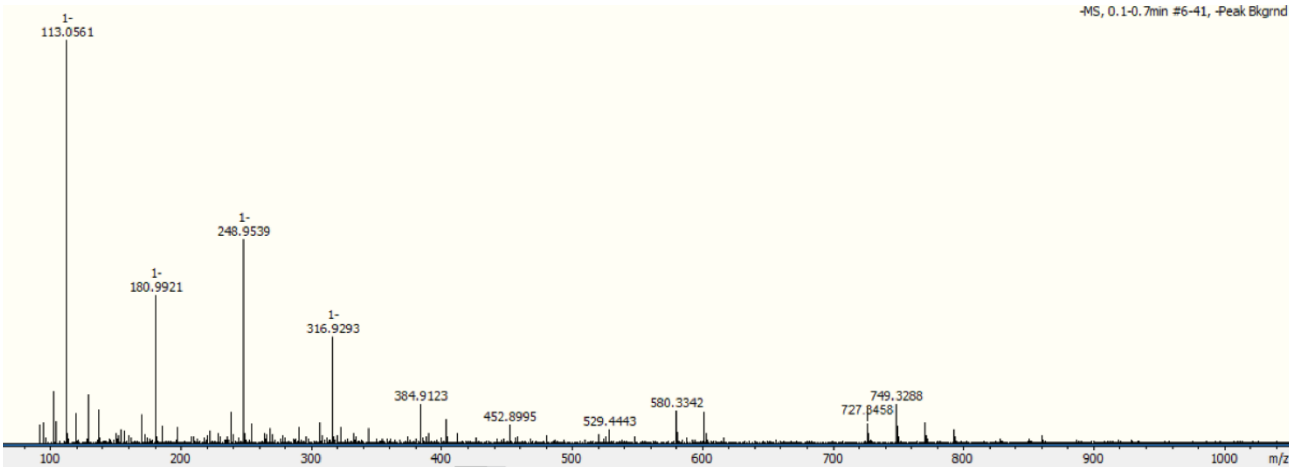

#### 1.6 HPLC traces and release experiment

Below is the HPLC traces of compound **10**, **11**, and **12**.

##### 1.6.1 HPLC trace of Compound 10

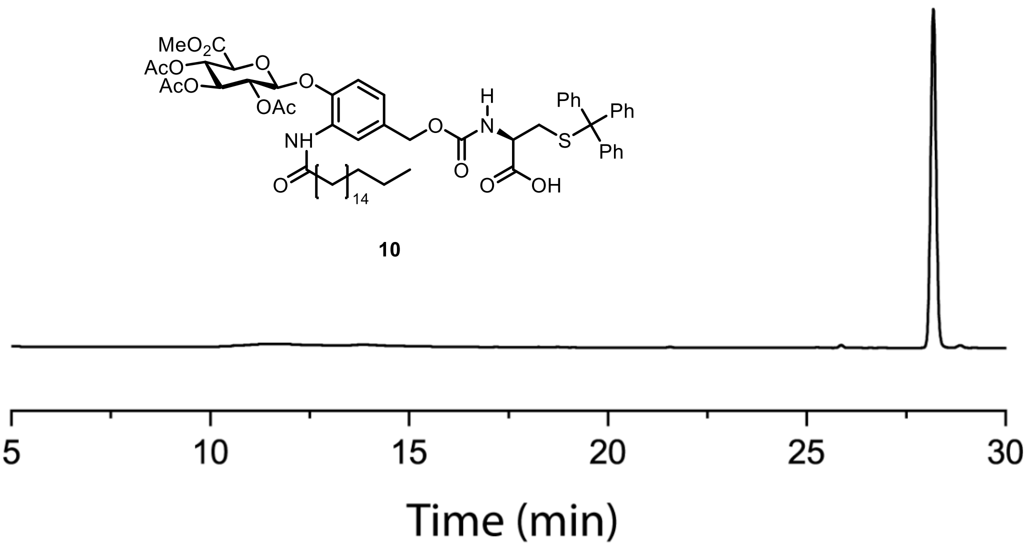

##### 1.6.2 HPLC trace of Compound 11

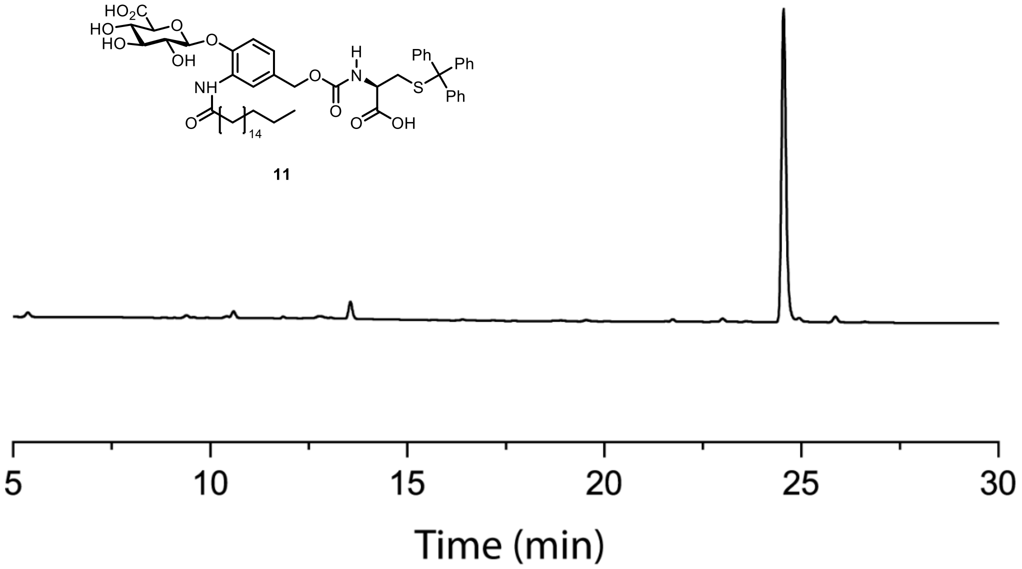

##### 1.6.3 HPLC trace of Compound 12

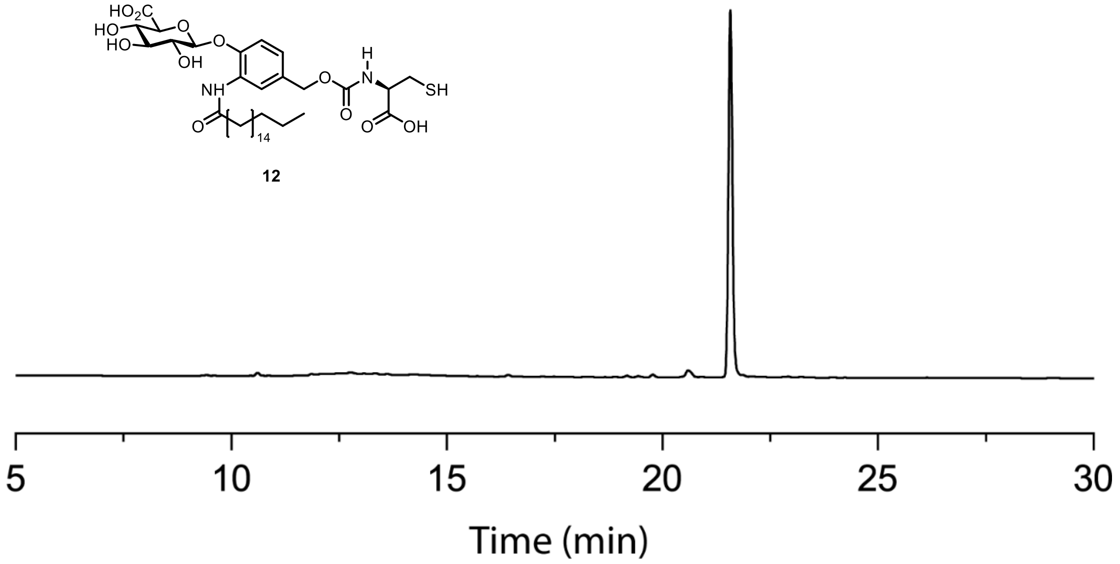

##### 1.6.4 HPLC release experiment

To prove the successful degradation of compound **11** when treated with the β-glucuronidase (GUS) enzyme, a HPLC release study was carried out. The release study was performed on compound **11** instead of the final receptor, compound **12**, to be able to detect by UV absorbance using the trityl group and to avoid oxidation of the thiol. The experiment was performed as described below:

A solution of compound **11** was prepared in PBS buffer (10 mM, pH = 7.4) with 4% DMSO at a concentration of 0.1 mM. The compound was incubated for 2 hours at 37°C either in the absence or the presence of the GUS enzyme (0.1 g/L). A reference with trityl cysteine was prepared in a similar manner. The GUS-treated sample was spin-filtered through a 30K spin-filter to remove the enzyme and injected into the HPLC using the standard method described in general information. The result of the degradation study is shown in Figure 2B.

### 2. Biological evaluation

#### 2.1 General information

FITC-casein was purchased from ThermoFisher. Kinetic experiments were performed on an EnSpire Multimode Plate Reader. Dynamic light scattering (DLS) experiments were performed on a Malvern Zetasizer Nano S90 and performed on all liposome samples to ensure successful extrusion and that the liposomes maintained their composition upon treatment with the EAR. Fluorescent microscopy was performed on a Zeiss Axio Observer Z1. Confocal laser scanning microscopy was performed on a Zeiss AixoObserver LSM 700.

##### 2.1.1 Buffer preparation

The buffer used for all experiments was a 20 mM PBS buffer with 1 mM EDTA and at pH 6.7. The buffer was prepared as follows: A solution of sodium phosphate dibasic in ultrapure water (1 M) and a solution of sodium phosphate monobasic in ultrapure water (1 M) were prepared. The two solutions were mixed while adjusting the pH to 6.7. The resulting mixture was diluted to give a final concentration of 20 mM. EDTA (1 mM) was added to the PBS buffer and the buffer was filtered through a 0.2 μm filter.

##### 2.1.2 Size-exclusion chromatography

The liposomes were after hydration, extrusion, and saponin treatment purified by size-exclusion chromatography (SEC). This was done by either a prepacked Sephadex® G-25 NAP-25 column from GE Healthcare Life Sciences, which was used for 800 nm liposomes (used for visualization) or a column packed with Sepharose® CL-2B, which was performed on 200 nm liposomes (kinetic studies). In both situations, the fraction size was 500 μL and DLS was used to identify the fractions with the liposomes.

#### 2.2 Experimental protocols

##### 2.2.1 Preparation of liposomes with encapsulated papain and papain substrates

A stock solution of the lipid alfa-phosphatidylcholine from egg yolk in chloroform (25 g/L) was prepared and 250 μL of that solution was added to a glass flask and cholesterol (2.09 mg) was further added to the flask. A lipid film was prepared in the flask by evaporation of chloroform using nitrogen flow while rotating the flask. The lipid film was further dried under vacuum overnight.

A solution of inactive papain (10 g/L, stock: 28 g/L) and the wanted substrate either $N_\alpha$-benzoyl-L-arginine-7-amido-4-methylcoumarin (1 mM, stock: 81 mM) or FITC-casein (0.2 mM, stock: 2 mM) was prepared and filled to 100 μL with buffer (20 mM PBS, 1 mM EDTA, pH = 6.7). The solution of papain and substrate in buffer was added to the lipid film and the liposomes were hydrated by vortexing for 10 minutes. After successful hydration of the lipids, more buffer was added to a total volume of 250 μL. The liposome mixture was extruded 15 times through a 200 nm filter (or 800 nm for visualization experiment). The resulting liposome mixture was treated with saponin (0.1 g/L) for 3 hours before the liposomes were purified by size-exclusion chromatography (SEC) yielding the wanted liposomes with encapsulated papain and papain substrate. For large experiments, the preparation of liposomes was scaled to suit the amount of liposomes needed.

##### 2.2.2 EAR incorporation into liposomes

The liposomes were prepared as described in section 2.2.1. The EAR, compound **12**, (stock: 27.4 mM in DMSO) was directly added to the mixture of purified liposomes and was left for 30 minutes for the receptor to anchor to the membrane. A control without receptor was prepared by adding the same volume of DMSO to the liposomes.

##### 2.2.3 Papain activation with EAR in liposomes – kinetic studies

The EAR-containing liposomes were prepared as explained above with a final concentration of EAR at 10 µM. The potential of the receptor was evaluated as a kinetic study, where the liposomes were treated externally with the GUS enzyme and fluorescent output by turnover of $N_\alpha$-benzoyl-L-arginine-7-amido-4-methylcoumarin was measured by plate reader. The samples were always prepared in the same way in a 96-well plate and each sample in triplicates; 50 µL liposomes (with or without EAR), 10 µL GUS enzyme (15 µg/mL, stock: 150 µg/mL), 40 µL buffer to a final volume of 100 µL. The control without GUS enzyme was treated with 10 µL of extra buffer instead. The GUS enzyme was added last using a multichannel pipette and the plate was directly subjected to the kinetic measurement. The experiment was reproduced in three independent experiment with the same preparation of liposomes.

##### 2.2.4 Papain activation with EAR in liposomes – visualization

The liposomes were prepared as described in section 2.2.1 with the change that during the hydration step a higher concentration of the substrate $N_\alpha$-benzoyl-L-arginine-7-amido-4-methylcoumarin (10 mM, stock; 81 mM) was used and the extrusion was performed through a 800 nm membrane. After preparation of the liposomes, the liposome mixture was treated with receptor (500 µM, stock; 27.4) for 30 minutes to let the receptor associate with the membrane. The liposome mixture was then purified on a NAP-25 column. The resulting mixture of liposomes was used to prepare following samples; 67 µL liposomes with EAR, 10 µL GUS enzyme (15 µg/mL, stock; 150 µg/mL), and filled to 100 µL with buffer or 67 µL liposomes with EAR and filled to 100 µL with buffer. Another control was prepared without EAR present in the liposomes; 67 µL (without EAR), 10 µL GUS enzyme (15 µg/mL, stock; 150 µg/mL), and filled to 100 µL with buffer. The samples were incubated at 37°C for 2 hours before they were analyzed using fluorescent microscopy.

##### 2.2.5 Variation in EAR concentration – kinetic measurement

The effect of variation in EAR concentration was performed similar to the experiment described in section 2.2.3 with five different final concentrations of EAR (250, 50, 10, 2, and 0.4 µM). Each sample was prepared in triplicates and the experiment was reproduced three times using the same liposome preparation. The kinetic data of fluorescence intensity vs. time of each sample was pooled together and the linear section of the plot was fitted to a line by linear regression on the software Graphpad Prism. The rate of reaction was estimated as the slope of the curve and plotted as function of EAR concentration.

##### 2.2.6 Degradation of casein and variation in GUS concentration

Liposomes were prepared as described in section 2.2.1 and 2.2.2 using the FITC-casein substrate and the kinetic studies were conducted in the same way as in section 2.2.3 with the exception that different concentrations of GUS enzyme was added to the wells (1.5 µg/mL, 15 µg/mL, or 150 µg/mL). The fluorescent output indicating casein degradation was monitored as plate reader experiment. Each samples was prepared in triplicates and the experiment was reproduced three times using the same preparation of liposomes.

##### 2.2.7 Visualization in GUVs

0.25 µg of lipids containing egg yolk phosphatidyl choline and cholesterol at molar ratio of 60:40 were made into a thin film on top of an ITO glass and left at vacuum overnight. Afterwards samples were hydrated with 300 µL $H_2O$ containing 300 mM sucrose 1 mM PBS. Then GUVs were formed by electroformation between ITO glasses (Nanion Technologies Vesicle Prep Pro, Freq: 10.0 Hz, Amplitude: 3.0 V, Temperature 37° C) for 3 hours. GUVs were stored at 4° C and used within 2-3 days after formation.

10 µL of GUVs were suspended in 100 uL of PBS (1 mM, 300 mM glucose) and EAR at 100 µM and fluorescein maleimide at 100 nM were added. A sample was subsequently treated for 1h with β-glucuronidase at 15 µg/mL. Control samples without treatment with β-glucuronidase, bearing only EAR or treated only with fluorescein maleimide were also prepared. Vesicles were analyzed by confocal microscopy. Imaging settings were adjusted and selected with the EAR+/Mal+ sample and kept constant for the remaining samples.

##### 2.2.8 Critical micelle concentration (CMC)

The method was adapted from Kalayanasundaram *et al* (*J. Am. Chem. Soc*. 1977, 99, 2039-2045). To solutions of EAR in PBS was added pyrene from a stock solution of 10 mM in ethanol to a final concentration of 10 µM. Final concentration of EAR ranged from 50 to 250 µM. Fluorescence of samples were recorded between 350 and 450 nm with excitation at 334 nm in a plate reader (Biotek Synergy™ H1 microplate reader). Then the ratio of fluorescence at $\lambda = 383$ ($I_3$) and $\lambda = 372$ ($I_1$) was plotted as a function of concentration of EAR and used to estimate the CMC (Figure S2).

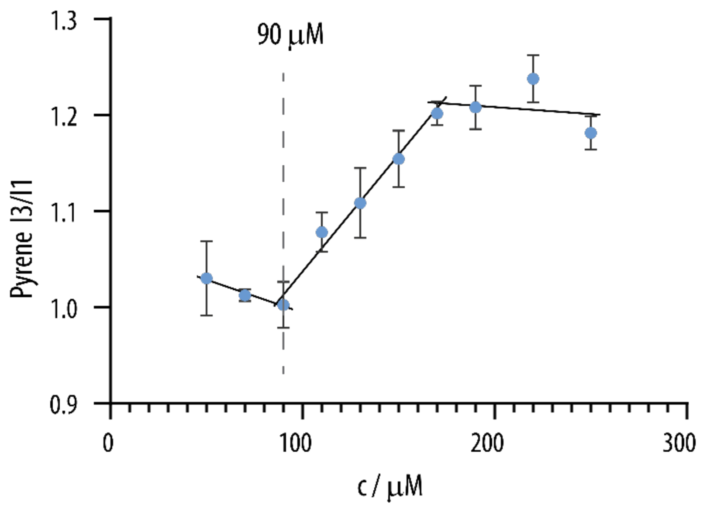

Ratio of pyrene $I_3/I_1$ fluorescence as function of EAR concentration in PBS.

##### 2.2.9 Comparison of papain activation in liposomes with cysteine and EAR

Liposomes were prepared as described in section 2.2.1 and 2.2.2. The kinetic experiment was performed similar to 2.2.3 with the exemption that the liposomes were diluted 5 times (10 µL was added to the plate instead of 50 µL). The liposomes were treated either with EAR (+/-GUS) or cysteine at matched concentrations of 2 µM.

**2.2.10 Inhibitor experiment in solution**

The effect of the lipid-impermeable maleimide 4-acetamido-4-maleimidylstilbene-2,2-disulfonic acid (AMS) reaction with free thiols (either cysteine or EAR) was first established as a solution experiment. This was to confirm successful reaction with thiols and thereby inhibition of papain activation. The experiment was conducted in a 96-well plate at a final volume of 100 µL with papain (0.03 g/L), fluorescein diacetate, FDA, as papain substrate (20 µM), and either cysteine or EAR at a matched concentration of 10 µM. The samples were treated with AMS (50 µM) for either 0 minutes or pre-treated for 30 minutes. A control sample was prepared without treatment with AMS. The evolution of fluorescence resulting from papain converting FDA to fluorescein was monitored as a kinetic experiment over 2 hours. All samples were prepared in three replicates and each experiment was further reproduced three times.

**2.2.11 Inhibitor experiment in liposomes**

The experiment in section 2.2.10 showed that the maleimide inhibitor, AMS, blocked the activation of papain by reacting with both cysteine and EAR in solution. Next, a similar experiment was conducted in liposomes. Papain-containing liposomes were prepared as previously described in section 2.2.1, however, without encapsulating a papain substrate. Instead, FDA was used as a membrane-permeable substrate. The liposomes were either treated with EAR (10 µM) or cysteine (10 µM) with and without the AMS inhibitor. The experiment was performed as a kinetic measurement of increase in fluorescence by the turnover of FDA to fluorescein. It was performed in a 96-well plate; 50 µL liposomes (with EAR or with cysteine), 10 µL GUS enzyme for EAR samples (15 µg/mL, stock: 150 µg/mL), 10 µL of AMS (50 µM, stock: 500 µM) and filled with buffer to a final volume of 100 µL. The samples were either treated for 0 minutes or 30 minutes with AMS. All samples were performed in three replicates and the experiment was reproduced three times using the same liposome preparation.

**2.2.12 Preparation of papain zymogen**

Papain was purchased from Sigma Aldrich as an inactive enzyme. Before use, this was established by measuring papain activity in solution with and without DTT (0.03 g/L papain, 5 µM papain substrate, and +/- 20 µM DTT). Typically the measurement confirmed papain inactivity and papain was used without further treatment. In case papain activity was observed, it was possible to block papain activity using following procedure.

Papain (5-10 g/L) and *S*-methyl methanethiosulfonate (500 equiv.) were dissolved in buffer (20 mM PBS, 1 mM EDTA, pH = 6.7) and left stirring overnight in the fridge. The resulting inactive papain was purified by Sephadex® G-25 NAP-5 column from GE Healthcare Life Sciences and finally concentrated by Amicon centrifugal filtration (MWCO 3 kDa).

**2.2.13 DLS Measurements**

DLS measurements were used to ensure that the liposomes had the wanted size and acceptable dispersities. All liposome samples were analyzed using DLS and the sizes were within the range of 200-250 nm and PDIs below 0.2. All measurements were conducted as three independent measurement. Figure S3 shows a standard DLS measurement result of 200 nm liposomes with and without EAR.

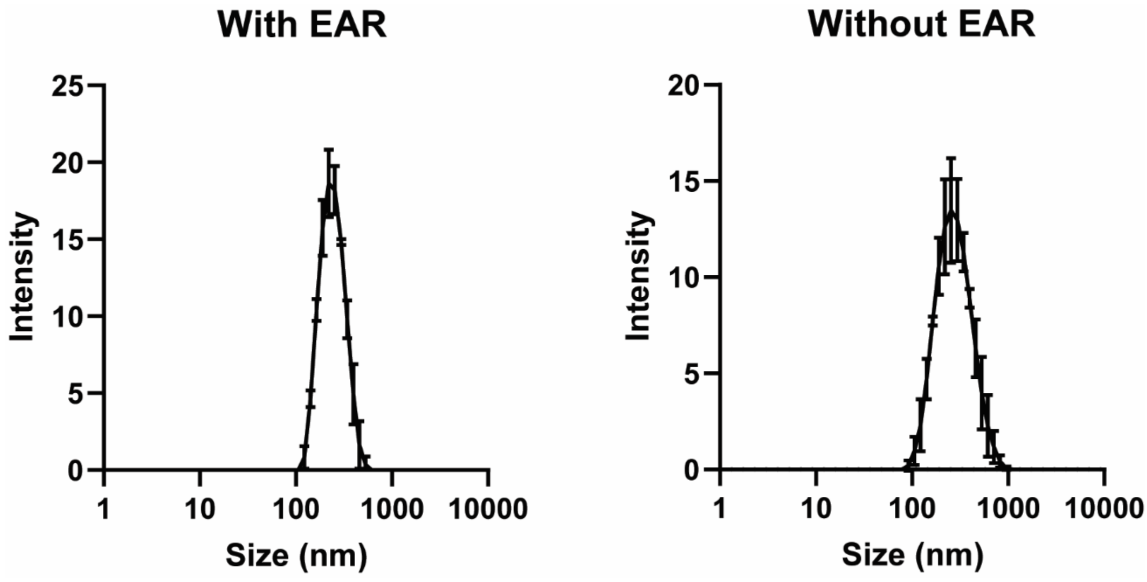

*Figure S3: DLS measurements of liposome samples (diluted 100 times) with and without EAR. With EAR; Z-average = 229 nm and PDI = 0.080. Without EAR; Z-average = 249 nm and PDI = 0.134.*

**2.2.14 Statistical analysis**

All statistical analyses are based on at least three independent experiments and are presented as mean ± SD. The statistical significance was established via either an unpaired t-test or a one-way ANOVA test with multiple comparison using the software Graphpad Prism, *** $p < 0.001$; n.s. = non-significant.
